# Anxious about rejection, avoidant of neglect: Infant marmosets tune their attachment based on individual caregiver’s parenting style

**DOI:** 10.1101/2023.05.18.541258

**Authors:** Saori Yano-Nashimoto, Anna Truzzi, Kazutaka Shinozuka, Ayako Murayama, Takuma Kurachi, Keiko Moriya-Ito, Hironobu Tokuno, Eri Miyazawa, Gianluca Esposito, Hideyuki Okano, Katsuki Nakamura, Atsuko Saito, Kumi O. Kuroda

## Abstract

Children’s physical, cognitive, and emotional maturation require adequate interactions and secure attachment with their primary caregivers. However, the causal relationships between parenting components and infant attachment behaviors remain unclear. New World monkey common marmosets provide an ideal model of human parent-infant relations because, like humans, infant attachment is shared among family caregivers, including parents and older siblings using intricate vocal communications. Combining the natural variations in parenting styles and subsecond-scale microanalyses of dyadic vocal and physical interactions, we demonstrate that marmoset infants signal their need by context-dependent call use and selectively maintain proximity with familiar caregivers. The infant attachment behaviors are individually tuned according to the caregiver’s parenting style; infants keep producing isolation calls when they are carried by rejecting caregivers and exhibit avoidance toward neglectful and rejecting caregivers. Family-deprived infants failed to develop such adaptive uses of attachment behaviors or age-appropriate autonomy and exhibited disorganized attachment behaviors. Thus, marmoset infants shape their ability to modulate their intricate attachment system depending on the caregiver’s attitude through early social interactions with their family caregivers. Demonstrating the significant similarity of the infant attachment system between marmosets and humans, these findings will pave a path to elucidating the neural mechanisms of the infant attachment system.

## Introduction

Early life adversity affects infants’ cognitive, social, and emotional development, ultimately increasing the risks of various medical conditions and premature death ^1–3^. Among the factors comprising the early life environment, the stability and quality of the relationship with the primary caregiver are critical for infants’ sense of security because infants are born immature and require extensive care for survival among all mammals. The experiences gained through interactions with the primary caregiver(s) (often the mother and other family members) are also essential for learning life skills and social behaviors in many species. Thus, infants have an innate motivation to seek and maintain proximity with the primary caregiver by a physical approach and signaling their need, collectively called the attachment system ^4^.

While the basic attachment system is innate, infants adjust their attachment pattern depending on the quantity and quality of care received. In humans, an appropriate caregiver is supposed to function as a *safe haven*, to which the infant returns for comfort and support, and as a *secure base*, from which the infant restarts exploration in the environment with a sense of security ^5^. It is theorized that if the caregiver inflicts fear or is insensitive to infant distress, infant attachment becomes “insecure”, i.e., infants are not fully confident in the caregivers’ availability and responsiveness ^6–8^. However, the discerned association between quality of rearing and attachment security is not large, possibly due to difficulties in controlling for other parameters such as genetic factors and the role of multiple caregivers in human studies ^9–12^. Thus, a nonhuman animal model should be established further to dissect the developmental mechanisms of infant attachment security. For this purpose, rodents have been studied extensively ^2,13,14^. Still, the attachment behaviors of rat or mouse pups are not as selective to a particular individual as humans, presumably because these species engage in communal nursing ^15^. In addition, the impacts of isolation rearing on the development of attachment behaviors or future social behaviors were less severe than the drastic effects in primates ^16–18^. In Old World monkeys, maternal separation or deprivation has been established to affect infant physiology, stress responses, cognition, and later social behaviors ^19–22^, although direct examinations of infant attachment security have been limited (see ^23,24^). Moreover, the mother is the sole primary caregiver in Old World monkeys, unlike humans ^25,26^.

Biparental or cooperative infant care in primates is limited to several family-living species, including New World monkeys Callitrichidae (marmosets and tamarins), *Plecturocebus* (titi monkeys), and *Aotus* (owl monkeys) ^27–31,32,33^. Among these, common marmosets (*Callithrix jacchus*) are a promising primate model with cutting-edge research resources such as genetic manipulation tools and multiple kinds of brain atlases and databases ^34–36^. At one birth, two infants are generally raised and carried almost continuously during the first postnatal months. Infant carrying impedes the carrier’s locomotor activities and thus is shared by the family members: the mother, father, and older siblings ^37^. Our previous study ^38^ identified two independent (allo)parenting parameters, *sensitivity* to infant distress and *tolerance* to infant carrying, similar to parenting styles established in humans ^39,40^. Furthermore, the molecularly defined subregion of the medial preoptic area in the basal forebrain specifically regulates caregivers’ *tolerance* in marmosets ^38^. These data suggest the common neurobiological mechanism of infant caregiving behaviors across primates.

Additionally, marmosets’ vocal development has attracted considerable attention because some of its characteristics are similar to those of human language, e.g., turn-taking and babbling ^41,42^. Undirected (isolated) vocalizations of infant marmosets undergo progressive changes from infant-specific “cry” and babbling to tonal “phee” calls. These changes have been attributed to the maturation of the vocal apparatus ^43,44^ and social learning from vocal feedback from parents ^45,46^. Nevertheless, as very young infants are continuously carried by the caregiver and infant calls function to attract parental approach ^47,48^, an investigation of infant call development within the context of parent-infant relations shall provide a fresh insight into vocal learning in marmosets ^49,50^.

Thus, this study investigates the relationship between parenting styles and infant attachment behaviors, including vocal communications, utilizing the high natural variations of the parenting parameters and the experimental manipulation of rearing conditions.

## Results

### The infant retrieval assay to study caregiver-infant interactions

The families were kept in a large family cage consisting of two to three connected cubicle cages. The infant retrieval assays, or “brief separation and reunion” from the infants’ viewpoint, were conducted utilizing two cubicle cages within their home cage. A caregiver, either a mother, father or an older sibling of the infant, was placed in one cage, and an infant in a wire basket was placed in an adjacent cage connected via a tunnel with a shutter (Fig. 1A) (Supplementary movie 1). After the shutter was opened, the caregiver typically entered the infant cage, approached, leaned into the basket, and came in contact with the infant. The infant climbed over the caregiver’s trunk, designated as infant retrieval and the start of carrying (Fig. 1B). After retrieval, the caregiver carried the infant for varying durations and may have eventually started rejecting the infant by rolling on the floor, pushing, and biting the infant. These behaviors often caused the removal of the infant from the carrier’s body (Supplementary movie 2). The session was continued for 600 sec after the first retrieval or from the assay start (see Table S1, Methods). Detailed analyses of 286 infant retrieval assay sessions were conducted involving 7 families, 25 infants, and 55 infant-caregiver dyads from postnatal day (PND) 1 to 36 (Table S1), and confirmed for sufficient inter-observational reliability with the on-site behavioral coding presented in our previous study (Fig. S1) ^38^. The caregiver’s and infant’s behaviors and calls listed in Table S2 were analyzed at a subsecond scale, with 0.2-sec bins using video and vocal recordings of the sessions. Based on the caregiver-infant interactions, the total period of each assay was broken down into five social contexts, which were mutually exclusive and collectively exhaustive (Fig. 1B): *Alone_BeforeRET,* when the infant was not carried yet before the first retrieval; *Holding,* when the caregiver was carrying the infant without locomotion; *Transport,* when the caregiver was carrying the infant and locomoting; *During_Rejection,* the period from the start of rejection to 9.4 seconds after the end of rejection (see below and Fig. 1B legend for this definition); and *Alone_AfterRET,* when the infant was not carried after the first retrieval occurred (Table S2).

**Fig. 1.**
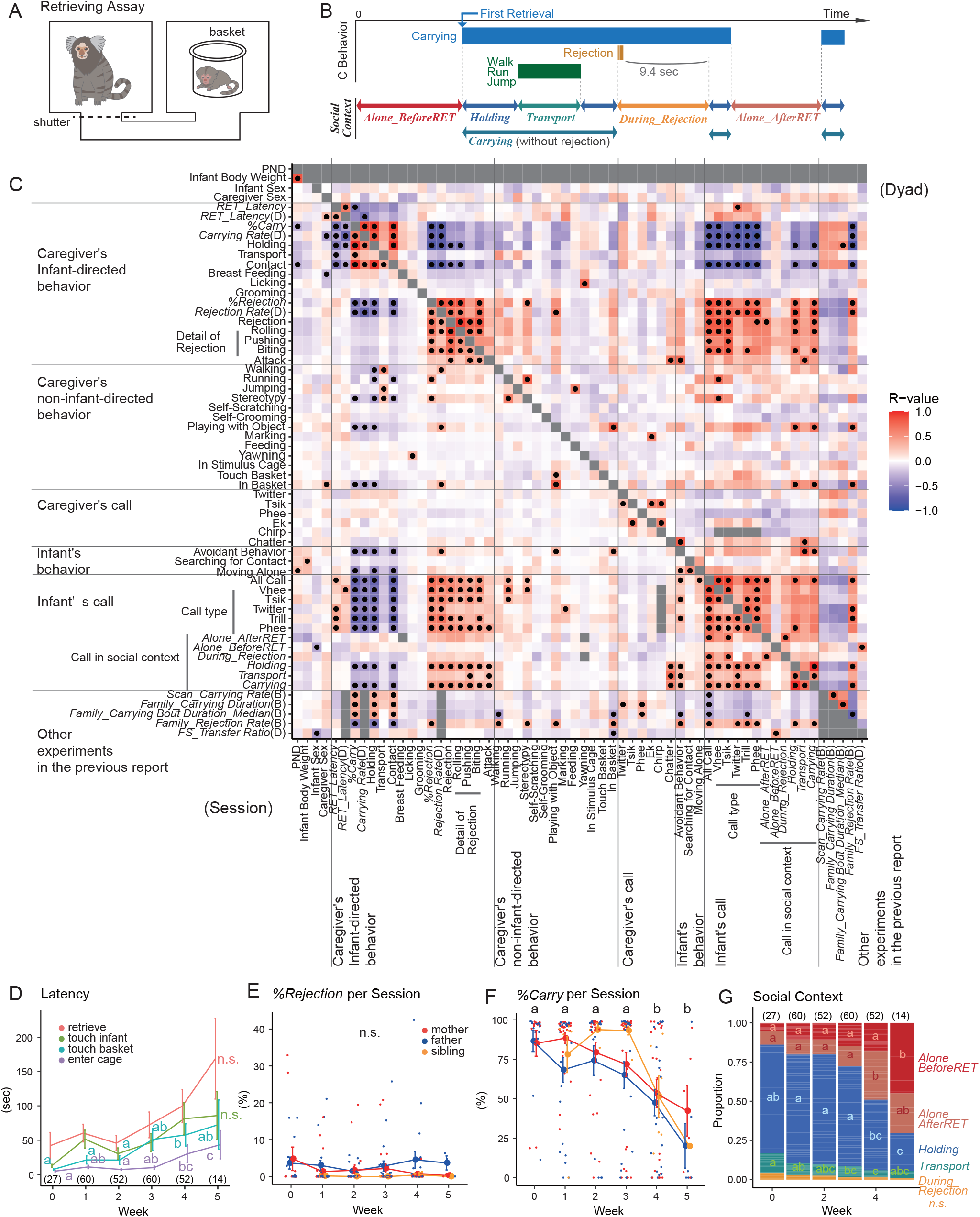
Caregiver behavior in retrieval assays. A Schematic of the retrieval assay in the home cage. B Caregiver-infant interactions and the five social contexts for the infants. Carrying includes transport and holding. A 9.4-second offset (see Fig. 2B) in *During_Rejection* was set to include the infant behaviors directly under the influence of the preceding rejection. C Correlation matrix of the parameters observed in the retrieval assays. The color indicates correlation coefficients (*r-*value, see Table S4). Filled circles, *p* < 0.05 (adjusted with Holm’s method, see Table S5). The left-bottom and right-top triangular parts show the parameters calculated per session and per the value averaged for each dyad, respectively. The bottom five parameters ending with either (B) or (D) are the parenting parameters derived from our previous study (Table S3) ^38^, which are averaged for each dyad (D) or each birth (B). D Mean ± standard error (s.e.) latencies of the caregiver’s behaviors after the shutter’s opening. Red: the first retrieval, yellow-green: the first touch of the infant, blue-green: the first touch of the basket, purple: the first reach of the infant cage. E-F Mean ± s.e. *%Rejection* (E) and *%Carrying* (F) in each postnatal week (filled circles and error bars). Red: mothers, blue: fathers, yellow: older siblings. Dots show individual sessions. G Proportion of duration of each social context. Red: Alone before the first retrieval, pink: alone after the first retrieval, blue: holding (carrying the infant without caregiver locomotion), green: transport (carrying the infant with caregiver walking, running, or jumping), yellow: during rejection. For D-G, different letters indicate significant differences among weeks (GLMM, *p* < 0.05), and the numbers within parentheses are the numbers of the sessions.

### Caregiving parameters define parenting styles

We first examined the Pearson’s product-moment correlation matrix of all the parameters of caregivers’ and infants’ behaviors (Table S2, 3) during postnatal weeks 0-4, to screen correlations between any parameters within the caregiver-infant dyadic relationship (Fig. 1C, Table S4, 5). We included the parameters representing the dyadic relations in other social contexts, such as undisturbed family observations and food-sharing experiments obtained in our previous study (Table S2, 3) ^38^. The significant correlations were screened from the left-bottom half of Fig. 1C for parameters derived from each infant retrieval assay session, or from the right-top half for parameters constant or averaged across each caregiver-infant dyad, and those candidate correlations were further analyzed below.

The screening analysis shown in Fig. 1C indicates that infant behaviors have high correlations with the caregiver’s infant-directed physical behaviors. Among these, we defined three caregivers’ parameters: (i) the <u>retrieval latency</u> (*RET_Latency*), the time required for infant retrieval, which negatively represented the *sensitivity* of the caregiver toward infant distress vocalizations; (ii) the <u>rejection rate</u> (*%Rejection*), which negatively represented the *tolerance* of the caregiver to infant carrying; and (iii) the <u>carrying rate</u> (*%Carry*), the total carrying duration divided by the session length, which represented the total infant care quantity. *RET_Latency* and *%Rejection* were mutually independent (*r* = 0.2338, *p* > 0.05), and both contributed to net infant carrying care (*%Carry – RET_Latency*: *r* = −0.5946, *p* < 0.001, *%Carry - %Rejection*: *r* = - 0.5969, *p* < 0.001). These caregiving parameters were highly correlated with the caregiving parameters in family settings (e.g., *FamilyRejectionRate(B)*) ^38^, indicating the existence of a caregiver-inherent “(allo)parenting style”.

Along with infant development, the latencies of caregivers’ approach to the infant increased (Fig. 1D), while *%Rejection* remained consistent (Fig. 1E). The net *%Carry* (Fig. 1F; *r* = - 0.2845, *p* = 0.0067) and the caregiver-infant contact declined after postnatal week 4 (*r* = −0.2808, *p* = 0.0094), indicating that the parent-infant interactions were essentially stable during the first postnatal week. No significant differences among mothers, fathers, or older siblings were found in these core caregiving parameters (Fig. 1E, F), while male caregivers were found to exhibit more “in basket” (a kind of object play) and naturally, shorter periods of breastfeeding (the left-bottom of Fig. 1C, *r* = 0.3295, *p* < 0.001 (in basket); *r* = −0.2643, *p* = 0.0331 (breastfeeding)).

Compared to the caregivers’ infant-directed behaviors, their vocalizations (calls) showed less-pronounced associations with infant’s behaviors (Fig. 1c), with only exception of caregiver’s chatter, which typically represents anger, correlated significantly with infant calls during carrying. Thus we focus on the caregiver’s infant-directed behaviors hereafter.

### “Approach” components of the attachment system of infant marmosets

Infants seek and maintain the proximity of specific familiar individuals using two kinds of attachment behaviors: 1) *approach* (seeking, following, clinging) and 2) *signaling* (crying, smiling and gestures) ^4^. In the first postnatal month of marmoset infants, “approach” behaviors were manifested mainly by actively clinging to the caregiver’s body whenever the caregiver made contact and by not breaking the contact unless the caregiver rejected them. *Infant_Moving_Alone* (Table S2) or spontaneous locomotor activities were positively correlated with infant age (*r* = 0.2979, *p* = 0.0023; the left-bottom half of Fig. 1C).

To examine the *selectivity of infant approach* behavior, we compared the speed of infants clinging to their parents or unfamiliar adults with multiple parental experiences (Fig. 2A). Infants instantaneously clung to their parents but not to the unfamiliar caregivers (Fig. 2A, left). In contrast, parental latencies to contact with their own or unfamiliar infants were highly variable and did not reach statistical significance (Fig. 2A, middle, the generalized linear mixed model (GLMM), p = 0.2040). The carrying rate, which both infants and parents could contribute to, was low for the unfamiliar parent-infant dyad (Fig. 2A, right). This selective attachment toward their own parents was attributed to familiarity rather than biological relatedness because fostering early neonates is generally successful in marmosets ^51^. Taken together, these results strongly suggest that marmoset infants develop a selective attachment with their family caregivers.

**Fig. 2.**
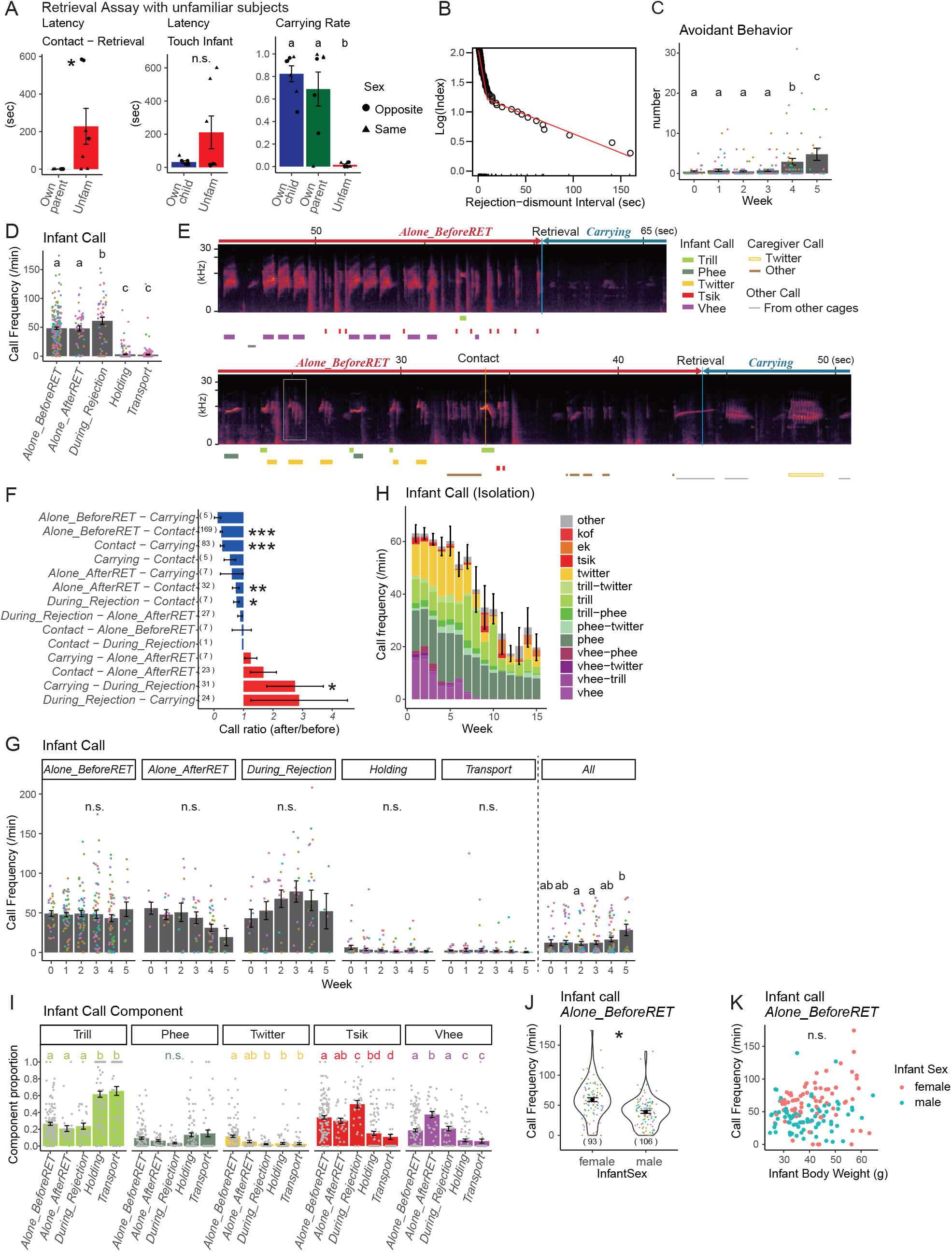
Infant behavior in retrieval assays. A Retrieval assay using unfamiliar (Unfam) or familiar caregivers (own parent or own infant). B Log-survivorship analysis of the rejection-dismount interval, defining the dismounts that occurred later than 9.4 seconds after the end of rejection as voluntary dismounts. Red line: the segmented regression line. C Mean ± s.e. numbers of avoidant behaviors per session. D Mean ± s.e. frequencies of infant calls in each social context. E Two typical spectrograms of infant and caregiver call at the transition from *Alone_BreforeRET* to *Carrying*. After the first contact and retrieval, the infant calls immediately stopped. The spectrogram of the infant twitter marked with the gray box is shown in Figure S5D. F Mean ± s.e. changes of the call frequency 10 sec before/after the shift of the social contexts (i.e., ratio = 1 means no change). Wilcoxon signed rank test with continuity correction, *: *p* < 0.05, **: *p* < 0.01, ***: *p* <0.001. The numbers within parentheses are the numbers of the scenes analyzed. G Mean ± s.e. call frequencies in each social context. H Mean ± s.e. total call frequencies and the composition of infant call types in the isolation recording (n = 9, four males and five females). I Mean ± s.e. call proportions over the social contexts. J Vaiolin plot of the call frequency of male and female infants during *Alone_BreforeRET*. The numbers within parentheses are the numbers of the trials. K Scatter plot of body weight and infants’ call frequency during *Alone_BreforeRET*. The call frequency was not affected by body weight (GLMM). Female: orange, male: green. L For A, C, D, G, I, and J, the asterisk and different letters indicate statistical significance at *p* < 0.05 in the GLMM. For C, D, J, and G, the dots show the values of each session, and the colors indicate the caregiver.

*Contact-breaking* behavior by infants was seen in two ways: one was when the infants were in contact with a caregiver but did not cling (passive), and the other was voluntary dismounting from the caregiver’s body (active). Practically, however, it was not straightforward to determine whether an infant’s dismounting was forced by caregivers’ rejection or by infants’ voluntary action. To empirically determine the direct influence of the preceding rejection on infant dismounting, we performed a segmented regression analysis to identify an abrupt change in the response function of a varying influential factor ^52^. The result in Fig. 2B identified the inflection point at 9.4 sec; thus, we defined *During_Rejection* as the period from the onset of rejection to 9.4 seconds after the end of rejection (Fig. 1B). The dismounting that occurred after a 9.4-second offset or those without preceding rejection were regarded as voluntary.

An infant’s *avoidant behaviors* were defined as the sum of voluntary dismounting and absence of clinging when the infant was in contact with the caregiver. The frequency of avoidant behaviors per session was substantially increased after postnatal week 4 (Fig. 2C), indicating that the marmosets become autonomous. Thus, these results suggest that avoidant behaviors after postnatal week 4 are a sign of typical development, while premature (i.e., within postnatal weeks 0-3) avoidant behaviors are atypical and may be a sign of decreased infant attachment to the given caregiver (see below).

### Infant call as a “signaling” component of the marmoset attachment system

#### Calls during rejection

Next, we studied infant vocalizations in various social contexts with the caregiver. The infants emitted calls most frequently when they were rejected or immediately after (Fig. 2D, *During_Rejection;* Supplementary movie 2, Fig. S2A). In the family observation, the infant’s vigorous calls during rejection appeared to attract other family members to the infant-carrier dyad and allow the infant to transfer quickly from the previous carrier to the next (Supplementary movie 3), suggesting the signaling function of infant calls. In this way, during the first three postnatal weeks, most infants directly transferred from one carrier to the next (see below for direct and indirect transfer of infants).

#### Calls while not being carried

Infants frequently vocalized also when they were not carried (Fig. 2D), often as a continuous string of various different call types termed “babbling” (Fig. 2E) ^41,49^. We observed that these intense bouts of calling stopped immediately after the infants were carried (Fig. 2D-E). Comparisons of call frequencies at the transition of social contexts revealed that infants reduced calling when they came into contact with the caregiver with any part of the body and further withheld calling when they clung to the caregiver’s body (= carrying) (Fig. 2F). These findings further support the notion that these bursts of infant calls in *Alone* contexts signaled the separation distress widely observed in mammalian infants ^53^ and thus were withheld immediately after contact with the caregiver.

#### Ontogeny of calls in family and isolation

The call frequencies during each social context did not change until postnatal week 5 (Fig. 2G). The increase in total calls during the test session along with infant development (Fig. 2G, right) should be attributed to the significant increase in *Alone* contexts (Fig. 1G). In a separate experiment, when we briefly isolated these infants and recorded their vocalizations in isolation, total call frequencies were high in the beginning and declined after postnatal week 6 (Fig. 2H), when parental carrying declined rapidly ^38^. These findings altogether suggest that the calls during *Alone* contexts (calls while not being carried in the family or in isolation) decline during infant development due to the decline in attachment needs by infant maturation (see Discussion).

#### Selective use of call types

Infant marmosets emit various kinds of call types, including twitter, tsik, trill, phee, and vhee [originally termed “ngâ” ^54^ or “cry” ^49^; here we use “vhee” to avoid unintended anthropomorphic interpretation] calls as well as various combinations of these calls (Fig. 2E, H, S2B, C). The ratio of infant call types depended considerably on the social context; infants emitted more trill calls when they were carried and more tsik calls when they were rejected (Fig. 2I). Twitter and vhee calls were most frequent during *Alone_BeforeRET* and *Alone_AfterRET*, respectively. These call type usages did not change substantially during postnatal weeks 0-5 in the infant retrieval assays (Fig. S2C). Thus, infant marmosets in the first postnatal month already use multiple call types selectively in each social context, although not exclusively.

In the isolated recording condition (Fig. S3A-G), while the total amount of phee calls remained stable across development (Fig. 2H, Fig. S3B), vhee and twitter calls rapidly declined during the first 8 weeks (Fig. 2H, Fig. S3C, F), which accounted for the decrease in total calls, indicating that the developmental increase of phee calls was relative and caused mainly by the developmental decline of total calls, especially vhee and twitter calls in isolated recording conditions.

As the only sex difference identified in infant attachment behaviors, female infants called more than male infants when the infants were alone before the first retrieval (the left-bottom half of Fig. 1C, 2J). This effect was not attributed to the sexual difference in body size growth, as there was no correlation between the isolation calls and body weight (Fig. 2K).

### Infants tune their attachment behaviors according to the parenting styles of each caregiver

We next investigated the relationship between parenting styles and infant behaviors by utilizing the wide range of individual variabilities in caregivers’ parameters (Fig. 3A-C). As an example, male twin infants Saku and Gaku (Fig. 3A) had a tolerant mother, Kachan, and an intolerant, rejecting father, Tochan, which occasionally attacked his infants. In their representative retrieval sessions, the mother (Fig. 3A, top panels) retrieved the infant quickly and carried it throughout without rejections. Both infants called only during *Alone_BeforeRET,* except for occasional trill and phee calls by Saku. The infants did not show any avoidant behaviors toward their mother, typical for this age. In contrast, the father (Fig. 3A, bottom panels) repeated retrieval and subsequent rejection, resulting in fragmented and short carrying bouts. In the father’s sessions, both infants exhibited premature avoidant behaviors (black triangles in Fig. 3A) not only during *During_Rejection* but also during *Carrying* (*Holding* + *Transport*). Moreover, the infants did not immediately withhold calling after the father’s retrieval (for example, approximately 600 sec in the left-bottom panel). These patterns of parental behaviors were consistent across litters (Fig. 3A, left and right), throughout infant development and across two births (Fig. 3B left, dark blue for Tochan, pink triangles for Kachan). These observations suggest that each parent behaved similarly toward different infants, and each infant behaved according to the caregiver’s behavior.

**Fig. 3.**
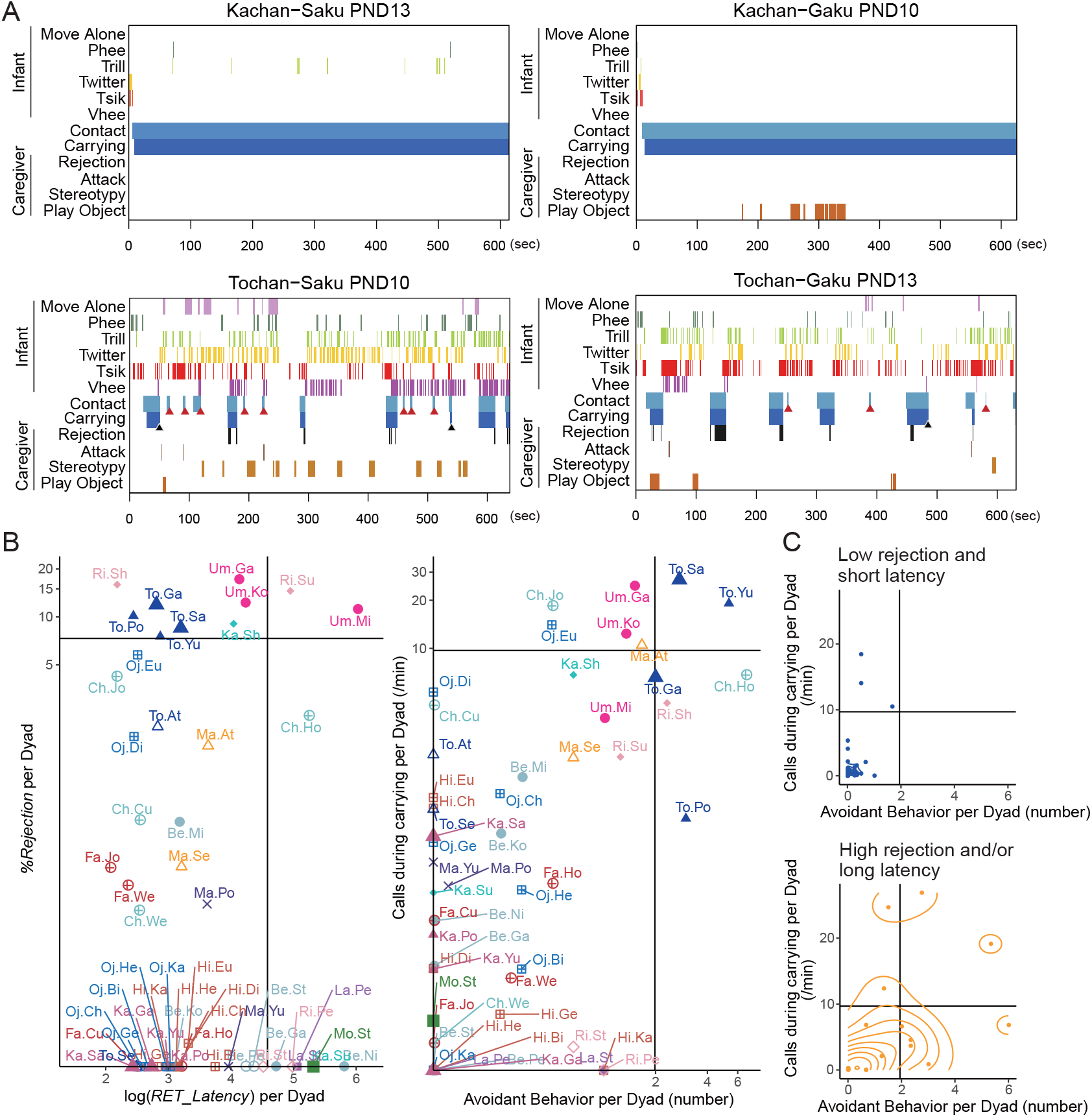
Relationship between caregiving parameters and infant behaviors. A Representative raster plots of the retrieval assay using a pair of littermates, Saku and Gaku (PND 10 and 13), by the mother Kachan and the father Tochan. Black triangles: dismounting without preceding rejection within 9.4 sec. Red triangles: refusal to cling when contacted. B Variation in caregiving (left) and attachment behavior (right) during postnatal weeks 0-3 (54 dyads, excluding one dyad in which the caregiver never retrieved the infant). Each point represents the value averaged for each caregiver-infant dyad, with the color of each caregiver (warm colors: females, cold colors: males). The marker shapes indicate each breeding pair. Labels near markers are the first two letters of the caregiver’s and infant’s names. The vertical and horizontal lines show the mean + 1 s.d. C Frequency of infant calls during carrying and the number of avoidant behaviors averaged for each dyad. Upper: both *%Rejection* and the log of *RET_Latency* were less than the mean + 1 s.d. (n = 36), lower: the other dyads (n = 18). Two-dimensional probability density was overlaid as a contour with 10 bins. Vertical and horizontal lines indicate the mean + 1 s.d. for each axis.

### Avoidance and anxious calls during carrying are associated with a low quantity and quality of caregiving

The above observations suggest that there are at least two kinds of atypical attachment behaviors of marmoset infants toward inappropriate caregiving: i) <u>avoidant behavior</u>, namely, dismounting and refusal to cling at contact, exhibited prematurely (i.e., within postnatal weeks 0-3) and voluntarily (i.e., not in *During_Rejection*); and ii) <u>calls while being carried</u>, namely, infant calls during the caregiver’s carrying. Indeed, the mappings of individual dyads for the average caregiving parameters (Fig. 3B, left) and the average atypical attachment behaviors (Fig. 3B, right) impressively resembled each other. The atypical attachment behaviors were much more frequent with the caregivers showing either *%Rejection* or *RET_Latency* over the mean + 1 S.D. (Fig. 3C, bottom) than the rest of the caregivers (Fig. 3C, top).

To examine the relations between parenting styles and infant attachment, we divided dyads into two groups, high/low groups for infant avoidant behavior and calls while being carried, and compared three caregivers’ parameters between the groups using the generalized linear mixed model (GLMM). The amount of infant avoidant behaviors was correlated with all the caregiving parameters *RET_Latency*, *%Rejection*, and *%Carry* (Fig. 4A-C, Fig. S2D-E), even when the caregivers’ behavior was equivalent at the moment of the infants’ responses. It suggests that infants avoid any caregivers showing insensitivity, intolerance or scarce carrying. On the other hand, the number of calls while being carried was associated with *%Rejection, %Carry* but not with *RET_Latency* (Fig. 4D-F, Fig. S2D-E), suggesting that infants call more while being carried by rejecting (but not insensitive) caregivers. The call type-specific analysis revealed that during carrying by rejecting caregivers, negative calls (tsik and vhee) were emitted more frequently, and the positive trill call was emitted less (Fig. 4G). Considering the fact that tsik and vhee calls were frequently emitted during isolation, and trill calls were emitted while being carried (Fig. 2I), infants might be insecure and exhibit anxiety-like calls when they were carried by rejecting caregivers. These atypical attachment behaviors were positively correlated with each other (Fig. 4H).

**Fig. 4.**
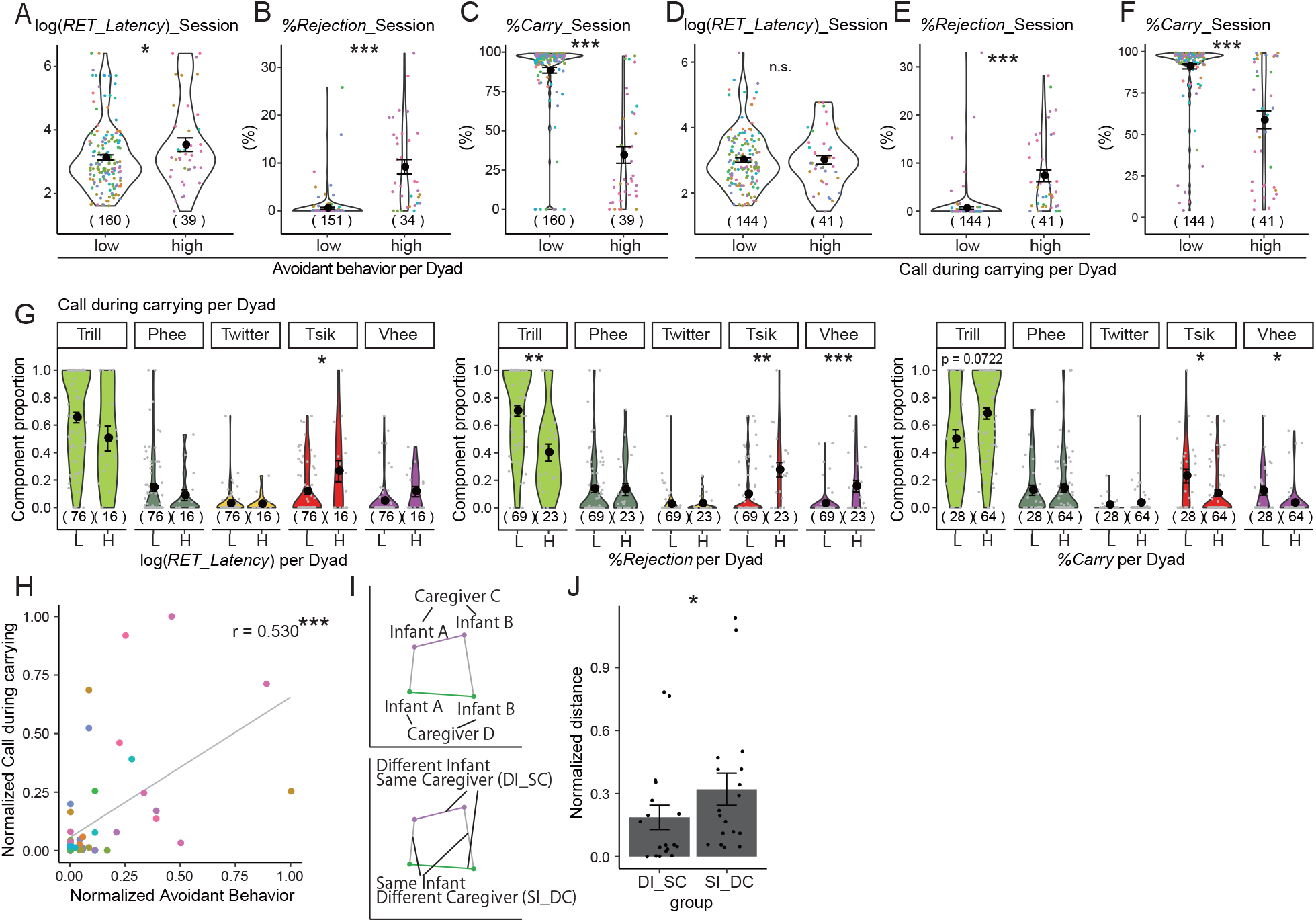
Infants tune their attachment behaviors according to the parenting styles of each caregiver. A-F Violin plots of the parenting parameters (A, D: *RET_Latency*, B, E: *%Rejection*, C, F: *%Carry*) in the groups of dyads with high/low infant avoidance (A-C) or high/low infant calls during carrying (D-F), defined by the average value. In B and E, 14 sessions without first retrieval were excluded. G Violin plots of the proportion of infant call types during carrying in each session in the groups of dyads divided by high (H)/low (L) *RET_Latency*, *%Rejection*, and *%Carry* at the average value. H Correlation between infant calls during carrying and avoidant behaviors. The Fig. 3B data are normalized to 0 to 1 by min-max normalization ((X – min(X)/(max(X) – min(X)) for both axes. The same color indicates data from the same caregiver. The gray line exhibits a regression line. *r* = 0.5301, *p* < 0.001, Pearson’s product-moment correlation coefficient. I Schema showing the method to compare the variability of two attachment parameters (H) of the same caregiver with two littermate infants (DI_SC, colored segments) and that of the same infant with two caregivers (either parents or older siblings) (SI_DC, gray segments). J Mean ± s.e. lengths of DI_SC (n = 18) and SI_DC (n = 18). Wilcoxon rank sum test, *p* = 0.0480. Each dot represents a distance. In A-G, the numbers within parentheses are the numbers of the sessions during postnatal weeks 0-3. GLMM, \**p* < 0.05, \*\**p* < 0.01, \*\*\**p* < 0.001. The black circles and error bars show the mean ± s.e. The dots show the values of each session, and the dot colors in A-F indicate the caregiver.

These correlations, however, do not infer the causality between caregivers’ and infants’ behaviors, and it is still possible that the atypical infant behaviors affect caregiving behaviors. We next compared the variabilities of atypical attachment behaviors within the same infant toward different caregivers (SI-DC, gray segments in Fig. 4I) vs. variabilities within the different infants toward the same caregiver (DI-SC, colored lines). We found that the variability between different caregivers and the same infant was larger (Fig. 4J). Together with our previous analysis showing the inherent nature of sensitivity and tolerance in each caregiver ^38^, this result suggests that the caregiver-infant relationship is determined more by consistent parenting styles of each caregiver than by infant predispositions.

### Family-separated, artificially reared infants as a model of highly insensitive caregiving: Study design and an example

The above-described data strongly suggest that parenting styles causally determine the pattern of infant attachment behaviors. Nevertheless, as these data are observational, the findings should be confirmed by interventional experiments (see ^22^). To this end, we utilized marmoset individuals who were separated from their families in infancy and reared artificially, modeling extremely insensitive caregiving. We collected 5 artificially reared infants (Art) from our breeding colony, which were born as triplets or to a mother with a dysfunction of one nipple (detailed in Table S6) and could not be maintained in the family even with supplemental formula feeding. Six of their littermates or age- and sex-matched infants were used as controls (Cont). These Art infants were housed individually and reunited with the original family at least 3 hours per day for 3-7 days per week in the daytime, except for one (Michael), which was rejected from the family in several trials of reunion and was once attacked (Table S6). The subjects were followed from birth and examined along with their development with family reunion observations and infant retrieval assays (Fig. 5A).

**Fig. 5.**
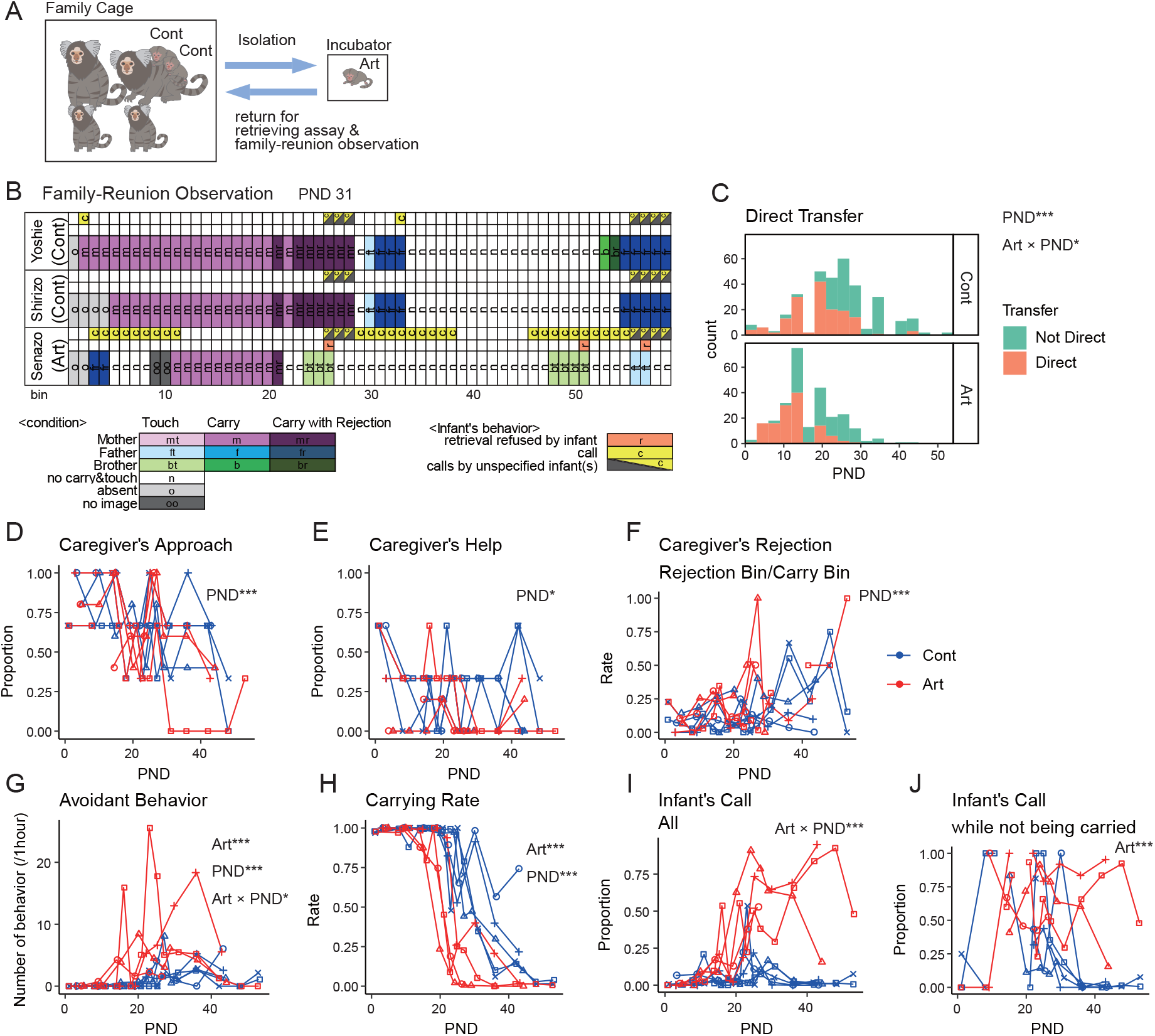
Artificially reared infants showed avoidance and age-disproportionate distress in family settings, even though the caregivers accepted them. A Schematic of the behavioral tests of Art and Cont infants. B Representative raster plot of family reunion observation. Bin: 10-sec. C Histograms of direct and nondirect transfers from one caregiver to another of Art (*N* = 4, 265 transfers) and Cont infants (*N* = 5, 324 transfers). Direct transfers decreased with infant PND (*z* = −7.50, *p* < 0.001) and declined earlier in Art (*z* = −2.27, *p* = 0.0232). D-J Caregivers’ (D-F) and infants’ (G-J) behaviors during the family reunion. Each marker shape represents each infant. Blue: Cont, red: Art. Numbers of sessions: 96 (D-I), 70 (J). Numbers of infants; 4 (Art) and 5 (Cont). *: *p* < 0.05, **: *p* < 0.01, ***: *p* < 0.001 (GLMM). D Proportion of the caregivers that approached the cage entrance upon infant return decreased with infant PND (*z* = −4.51, *p* < 0.001). E Proportion of caregivers that attempted to retrieve the returning infant by the forelimb or showed aggressive behaviors toward the experimenter decreased with PND (*z* = −2.43, *p* = 0.0149). F Caregiver’s rejection rate (rejecting bins / carrying bins) decreased with PND (*t* = 4.992, *p* < 0.001). G Infants’ avoidant behaviors was more frequent in Art (*z* = 4.80, *p* < 0.001) and increased with PND (*z* = 4.49, *p* < 0.001), and the increment was more pronounced in Art (*z* = −2.39, *p* = 0.0169). H Carrying rate (carried bins / total bins) was lower in Art (*t* = −5.20, *p* < 0.001) and decreased with PND (*t* = −15.29, *p* < 0.001). I Number of bins with infant calls increased with PND in Art (*t* = 5.30, *p* < 0.001). Number of bins with infant calls while not being carried was higher in Art (*t* = 4.45, *p* < 0.001).

*Family reunion* observations during infancy were performed with four Art and five Cont infants. The observation started by returning the infants to the home cage of their original family after body weight measurements, and the behaviors of the infants and the family members were coded in every 10-second bin. In a representative example (Fig. 5B), one Art and two Cont littermate infants at PND 31 were returned to their family cages one by one. The Art infant Senazo (the bottom row in Fig. 5B) was first retrieved by the father but soon dismounted because of the father’s rejection (dark blue bins). The experimenter assisted Senazo in clinging to the mother, which was tolerant and was carrying two Cont infants (Yoshie and Shirizo, the top and middle row of Fig. 5B). The mother carried three infants (10^th^ to 20^th^ pink bins) and then rejected them (21^st^ dark purple bin). Senazo dismounted, while two Cont littermates kept clinging even though the mother kept rejecting them for 6 consecutive bins. In the meantime, Senazo came into contact with the brother (24th-26th light green bins) but did not cling to the brother (retrieval refused by the infant, orange mark above the bin). Senazo called in two long bouts from the 29^th^ and 46th bins (yellow marks), and came into contact with the brother and the father sequentially but refused to cling. The two Cont infants did not call while being alone after dismounted from the father at the 34^th^ bin, which showed age-appropriate maturity (Fig. 2G). In contrast, Senazo showed excessive distress calls (which should attract caregivers) but avoided simultaneously, the behavior reminiscent of “disorganized” attachment in humans. In addition, compared to Cont littermates, Senazo tended to give up clinging and dismount easily by brief rejection.

### Artificially reared infants showed avoidance and age-disproportionate distress in family settings, even though the caregivers accepted them

The differences in the parent-infant relationship between Cont and Art infants during the family reunion observation were statistically examined in the GLMM, with explanatory variables of Art/Cont and PND (Fig. 5C-J). Typically, when an infant was returned to the home cage after a transient removal (e.g., for body weight examination), the caregivers often approached the returning infants, showed aggression to the experimenter whose hand grabbed the infant, and tried to reach and draw the infant closer by the forelimb. The proportion of caregivers showing this infant approach in the family declined significantly with infant age but was not different between Art and Cont infants (Fig. 5D). Subsequent caregivers’ attempts to retrieve the returned infant (caregivers’ help, Fig. 5E) and caregivers’ rejection (Fig. 5F) also did not differ between Art and Cont infants. Thus, marmoset caregivers appeared not to discriminate between Art and Cont infants.

On the other hand, infant avoidant behaviors toward caregivers were significantly more frequent among Art infants than among Cont infants after the second postnatal week (Fig. 5G). This might cause a decrease in the total carrying rate of Art infants, starting at a similar time (Fig. 5H).

The duration of each carrying bout, the latency for the first rejection in each carrying bout, and the total number of rejections during each carrying bout declined according to infants’ development (Fig. S4A-C). Among Art infants, the total number of rejections during carrying declined faster than among Cont infants (Fig. S4C), likely due to the reduced tolerance to rejection among Art infants (Supplementary movie 4). In concordance with this interpretation, the transfer of the infants from one caregiver to the next was different between Art and Cont infants (Fig. 5C). When rejected, Cont infants tended to stick to the caregiver until another caregiver came close and directly moved from the present caregiver to the next (direct transfer) until PND 20. However, Art infants tended to dismount to stay alone before being transferred to the next caregiver (nondirect transfer) after PND 12 (Fig. 5C). Overall, these data suggest that Art infants exhibited compromised approach components of attachment behaviors and increased avoidant behaviors toward caregivers, even though the caregivers behaved similarly toward Art and Cont infants.

Next, vocal behaviors were compared between Art and Cont infants (Fig. 5I, J). In the family reunion observations, the total calls as well as calls while not being carried were significantly more frequent among Art infants, particularly after PND 30 (Fig. 5B, I, J), indicating the age-disproportionate, excessive distress signaling of Art infants (note that in this family setting, calls while being carried could not have been assessed precisely and thus will be described below in the dyadic infant retrieval assays). We also assessed the development of vocalization of one Art infant Michael in the isolated recording condition (Fig. S3A-G). Compared with Cont infants (including Cont littermate Cubby), Michael exhibited fewer calls in the first postnatal month, possibly because Michael was used to staying alone, while Cont infants were not. After the second postnatal months, while Cont infants reduced the calls, especially vhee and tsik calls, rapidly, Michael did not show this trend, again consistent with the delayed independence compared to the family-reared infants.

### Avoidant attachment in Art infants in dyadic infant retrieval assays

We next conducted dyadic retrieval assays with four Art and four Cont infants (Fig. 6A). In an example case, Cubby (Cont) and Michael (Art) were tested with their father Chuck. While Cubby was carried throughout even during and after rejection (top panel, Fig. 6B), Michael clung on but dismounted from Chuck’s body soon after the first retrieval without obvious rejection from Chuck (left black triangle, Fig. 6B bottom) and then was carried again. Michael was rejected once, and a minute later, it dismounted from Chuck without preceding rejection. Chuck made contact again with Michael during 178-190 sec, but Michael did not cling onto Chuck (red triangle at approximately 190 sec, Supplementary movie 5) and kept calling. The refusal of clinging occurred again at the end of the session (red triangle, right). Overall, these patterns of atypical behaviors toward the caregiver were consistent with those observed during family reunion (Fig. 5B).

**Fig. 6.**
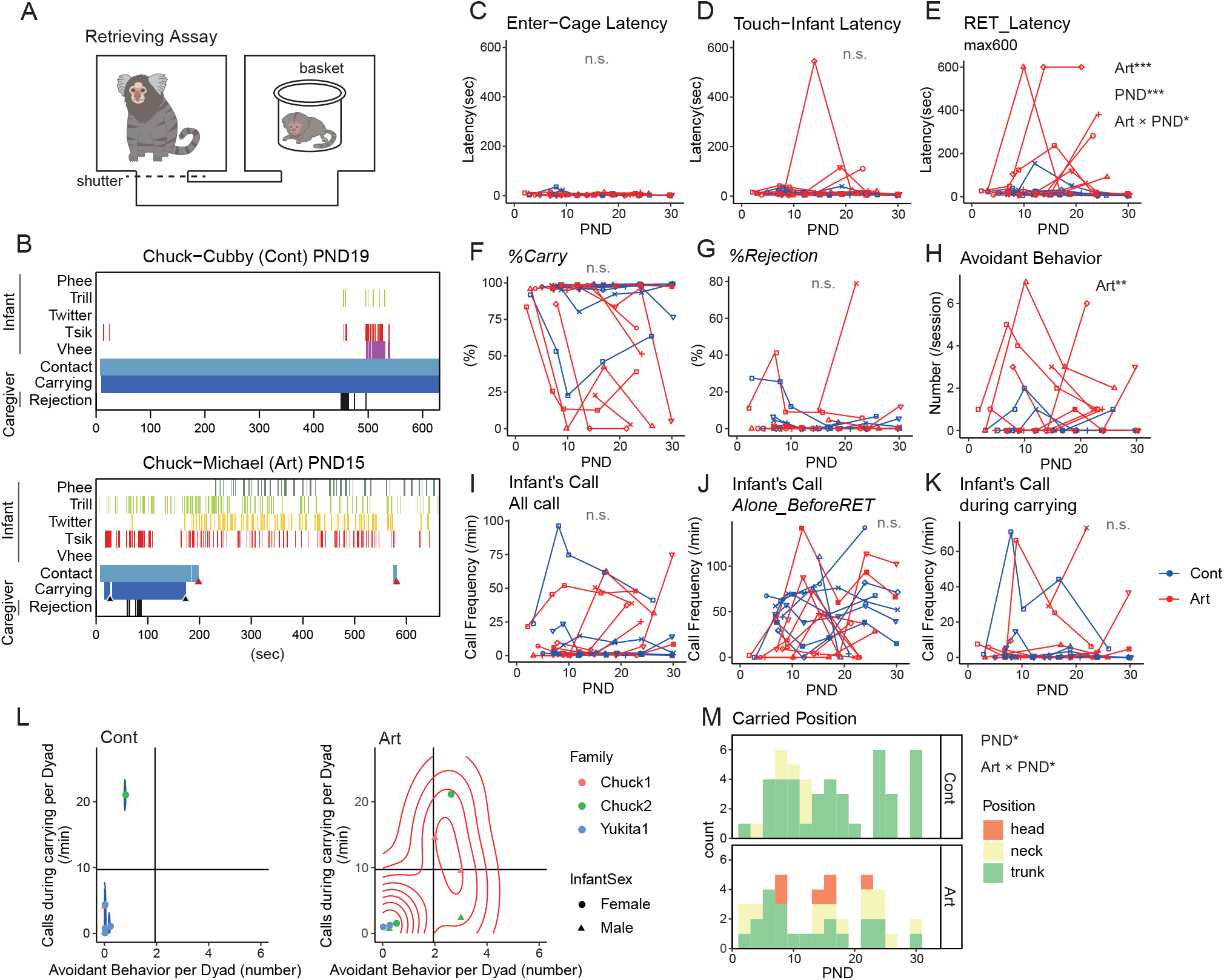
Avoidant attachment in Art infants in dyadic infant retrieval assays. A Schematic of the retrieval assay. B Two typical raster plots of the retrieval assay, performed by Cubby (Cont) and Michael (Art) and their father. Black arrowhead: voluntary dismounts. Red arrowhead: retrieval refused by the infant. C-K Caregiver-infant behaviors and interactions during the retrieval assay. The shape of the markers represents each dyad. Numbers of sessions: 69 (C-F, H-J), 66 (G, K; three sessions without retrieval were excluded). N of dyads =8 each. C Latency of caregiver’s reach to the infant cage. D Latency of caregiver’s touch to the infant. E Latency of infant retrieval, longer in Art (*t* = 3.61, *p* < 0.001) and the older infants (PND, *t* = 3.58, *p* < 0.001). The coefficient of PND was smaller in Art (*t* = −2.54, *p* = 0.0111). F Percentage of carrying. G Rejection rate. H Infants’ avoidant behaviors were more frequent in Art (*z* = 3.64, *p* < 0.001). I Frequency of infant calls. J Frequency of the infant’s call before the first retrieval. K Frequency of infant calls while being carried. L Infant calls during carrying and the number of avoidant behaviors in each dyad, overlaid with the two-dimensional probability density as a contour with 10 bins. Vertical and horizontal lines are the same as in Fig. 3B. The dot colors indicate the infants. The dot shapes indicate the infant’s sex. M Infant positioning on the caregiver’s body just after the retrieval (Cont: *N* = 4, 47 points, Art: *N* = 4, 40 points). Older Cont infants tended to hold onto the caregivers’ trunk (*z* = 2.01, *p* = 0.0449), while Art infants did less so (Art × PND: *z* = −2.12, *p* = 0.0342) and sometimes clung onto the caregiver’s head. *: *p* < 0.05, **: *p* < 0.01, ***: *p* < 0.001 (GLMM).

GLMM analyses revealed that the latencies of caregivers’ entry into the cage containing the infant (Fig. 6C) and contact with the infant (Fig. 6D), which solely depended on caregivers’ motivation, were not different between Art and Cont, while the actual retrieval latency was significantly longer for Art (Fig. 6E), possibly because of avoidance of Art infants. In the dyadic situation examined within PND 30, *%Carry* and *%Rejection* were not significantly different between Art and Cont (Fig. 6F, G). For infant behavioral parameters, infant avoidant behaviors were more frequent among Art infants (Fig. 6H), as in the family reunion observations. Infant total calls or calls during carrying did not differ between Art and Cont within PND 30 (Fig. 6I-K) (note that calls while not being carried were higher after PND 30 in family settings (Fig. 5J)). In the plot of atypical attachment behaviors for Cont and Art infants (Fig. 6L), Cont infants showed few avoidant behaviors and calls while being carried, the latter except for one dyad (Chuck-Nana; note that Chuck was inherently rejecting toward any infants as seen in Fig. 3B). In contrast, Art infants often exhibited more avoidant behaviors (Fig. 6L). Furthermore, Cont infants clung mainly to the caregiver’s trunk, especially when they became older (Fig. 6M, top, Supplementary movie 6). However, Art infants often clung onto the head or neck (Fig. 6M bottom, Supplementary movie 7). In summary, Art infants showed multiple atypical attachment behaviors, even when caregivers treated them equally to Cont infants in both family reunion and dyadic situations, indicating the pivotal role of social contact during the first two postnatal weeks in the proper development of the infant attachment system.

## Discussion

### Parenting styles in primate caregivers

This study investigated caregiver-infant interactions in common marmosets with high temporal resolutions and elaborated on caregiver-inherent (allo)parenting styles, comprising sensitivity, tolerance and caregiving quantity operationally defined in dyadic interactions. These parenting parameters showed significant consistency within each caregiver across infants and experimental paradigms. Furthermore, the sensitivity-tolerance dimensions of each caregiver had a substantial agreement with studies on maternal styles in Old World monkeys ^55–59^ and humans ^4,39,40,60,61^. Of course, human psychological constructs are much more complex than behavioral parameters in marmosets. Nevertheless, it is an intriguing consistency concerning the evolution of primate parenting and deserves further investigation.

### Typical and atypical infant attachment behaviors in response to individual parenting styles

This study established that isolated marmoset infants call vigorously and selectively attach to familiar caregivers. Upon contact with the caregiver, they immediately cling on and withhold calls. This clearcut on-off regulation of the infant vocalizations may solicit caregivers’ reinforcement learning to retrieve the infant, as loud infant calls should attract predators in the natural habitat. Both signaling and approach behaviors decline after one month of age, when the infants gain their locomotor independence. All these features of marmoset infant behaviors are in accordance with the core concept of the infant attachment system as defined by Bowlby ^4^.

This study next demonstrated that family-reared infants tune their attachment behaviors flexibly toward each caregiver, according to the caregiver-inherent (allo)parenting styles. Both rejecting and insensitive parenting increase premature avoidant behaviors in infants, even when other carriers are unavailable. These avoidant behaviors may be interpreted as the diminished *safe haven* quality of the caregiver-infant interaction, and may also be regarded as earlier independence earned as an adaptation to an adverse rearing environment (c.f., ^62^). Another atypical attachment identified here is the insufficient turning-off of vocalizations upon carrying by intolerant caregivers. Infants cling to the intolerant caregiver but simultaneously emit more negative calls (tsik and vhee calls) than trill calls (Fig. 4G), as if they were still alone. Calls while being carried may thus indicate anxiety and a decreased *sense of security*. And as the *sense of security* obtained in the proximity of the caregiver should be a prerequisite for the subsequent re-departure from the caregiver, calls while being carried may also be relevant to impaired *secure base* behavior.

It should be stressed here that these apparently “insecure” attachment behaviors of family-reared marmoset infants are not a fixed “style” or predisposition because infants selectively show secure or insecure attachment toward each caregiver (Fig. 3A). This finding points to the advantage of studying marmosets, in which infant attachment is shared among multiple caregivers, as in humans. While infants of macaque monkeys, dogs, and chicks have been reported to show unaltered or even stronger clinging when their attachment figures behave abusively (see ^63^), marmoset infants show evident avoidance in this study, possibly because they can still rely on other caregivers. Marmoset infants may be more resilient when one caregiver is abusive, while macaque infants receive direct effects from abusive mothers and show delayed independence and excessive anxiety ^23^.

### The effects of artificial rearing in marmoset infants and comparisons with disorganized attachment and other attachment-related issues in humans

In contrast to the flexibility of the attachment system in family-reared marmosets, artificially-reared infants exhibit rigid, fixated avoidance of any caregivers. Moreover, they do not develop age-appropriate autonomy and call excessively while not being carried after one month of age. These calls of Art infants do not function to gain proximity to the caregiver because they simultaneously show contact avoidance. These paradoxical behaviors of the Art marmosets bear some resemblance to the disorganized attachment ^64^ or the inhibited type of reactive attachment disorder ^65^ documented in humans, yet it is too preliminary to propose Art marmosets as a model of human attachment disturbances. Nevertheless, the present results highlight the pivotal role played by the care experienced in the first postnatal month (approximately the first year for human infants) in shaping both approach and signaling components of attachment behaviors, thus forming a basis for dissecting neuromolecular mechanisms of infant attachment and its dysregulations. We are currently studying whether and how Art infants’ atypical social and nonsocial behaviors extend into adulthood and which brain areas are involved in separation distress and reunion behaviors in young marmosets.

### Vocal development and the attachment system in marmoset infants

This study demonstrates that infant vocalizations are highly dependent on social contexts rather than on infant age during the first 6 weeks (Fig. 2G, I, S2C). Moreover, our analyses shows that the developmental shift from vhee to phee calls in isolated conditions is mainly caused by the rapid decrease of vhee and twitter calls in the first two months, resulting in the relative increase of phee calls, while the absolute amount of phee calls is stable during the postnatal 3 months (Fig. 2H, S3). Together with other findings in infant attachment behaviors, these observations collectively suggest that these developmental changes of infant calls under isolation can be at least partly caused by the developmental decrease in infant need for carrying. This notion is supported by the previous finding that even young adults emit vhee-like calls in a certain context, such as subordination ^54^. It is possible that vhee (cry) calls can be used for begging-like social intent of marmosets in general, and such situations decrease with infant maturation.

Chronic inhibition of naturalistic parent–infant interactions disturbs both early vocal learning and infant attachment formation; the latter hinders the maturation of autonomy and affect-regulation in primates. Both vocal learning and attachment formation can independently influence vocal patterns, depending on social contexts and developmental stages (e.g., Fig. S3A-G). As a result, these data are complex yet coherently understood utilizing the concept of the infant attachment system, which requires proper parental care to be matured ^19,22^.

We quickly add that the present findings neither contradict nor disregard the importance of vocal learning from adult feedback in marmosets. We also admit that our study is not elaborated or sufficient for vocal analyses, as our main focus is on parent-infant relations. The present study instead proposes the infant attachment system as an *additional* regulator of infant vocalization in various social contexts. Appreciating infant vocalization as a signaling component of the attachment system may help a precise understanding of the development of vocal patterns in infant marmosets, including babbling.

## Supporting information

Supplemental Talbes 1-8

## Acknowledgments

We thank Drs. Michael Numan and Joji Tsunada for fruitful discussion, Sayaka Shindo and Yumi Ogawa for marmoset breeding, and the RIKEN Center for Brain Science, Research Resources Division for animal care.

This research was supported by RIKEN Center for Brain Science (2014-2022) to K.O.K., the Japan Agency of Medical Research and Development (AMED) under grant numbers JP22dm0207001 to H.O. and JP20dm0107144 to K.O.K., Brain/MINDS project 2014 to K.O.K. and No. 16 dm0207003h0003 to K.N., and JSPS KAKENHI grant number JP26893327, JP16K19788, JP20K12587 to K.S., JP19K16901 to S.Y.-N., and JP18KT0036 and JP22K19486 to K.O.K.

## Author contributions

S.Y.-N. designed and carried out experiments described in Fig. 5-6, S3-6, analyzed data described in Fig 1-6 and S2-6 with help from T.K. and E.M. and produced the tables and figures. A.T. designed and carried out the microanalysis described in Fig 1-4, and S1-2 with support from G.E., S.Y.-N., K.S., and K.O.K. K.S. designed and carried out experiments described in Fig 1-4 and S1-2 with support from A.S. A.M. carried out a part of experiments described in Fig. 5, 6, and S4 with support from H.O. K.O.K. conceived of and organized the study with A.T., S.Y.-N., and A.S. and with support from K.M.I., H.T., and K.N., and wrote the manuscript with S.Y.-N., and with contributions from all the authors.

## Materials and Methods

### Animals (normative family)

All animal experimentation was approved by the Animal Experiment Judging Committee of RIKEN (equivalent of Institutional Animal Care and Use Committee, IACUC, approval numbers H28-2-210, H30-2-206, W2020-2-027) and was conducted in accordance with the 2011 guideline from the National Research Council of the national academies. Common marmosets were reared at the RIKEN Center for Brain Science in accordance with the institutional guidelines and under veterinarians’ supervision.

We tested 65 common marmosets from 9 families: 8 fathers, 9 mothers, 32 siblings (19 males and 13 females, 3 out of 32 were also included as two fathers and a mother later), and 49 infants (29 males and 20 females, 30 out of 49 were also included as siblings and/or fathers later). Detailed information about the subjects is shown in ^38^. They were housed as a family, ranging from parents and two infants (minimum) to parents, two older siblings, two younger siblings, and two infants (maximum). The average number of infants at one birth was 2.21. When triplets or quadruplets were born, the extra infant(s) were used for artificial rearing experiments or removed from the family at PND 0-2 to be subjected to histological experiments. In Chuck’s family, the mother (Fastener) was able to nurture only one infant at a time because of the dysfunction of one nipple. Thus, another infant was removed or, in one case, artificially reared (see below).

One cage was 43 (width) × 66 (height) × 60 (depth) cm. Two or three cages were joined through a square hole (9.6 cm wide × 10.5 cm high) on a side panel or a metal mesh tunnel (75 cm wide × 30.5 cm high × 21 cm deep) placed in front of two cages, depending on the number of family members in accordance with the ethical guideline of RIKEN, to form one single home cage. Each cage contained a food tray, a water faucet, two wooden perches, and a metal mesh loft. Although tactile contact was restricted between families, visual, olfactory, and auditory communication was possible in the colony room. Water and food were supplied ad libitum. The monkeys’ food was replenished at approximately 11:30, and supplementary foods, such as a piece of a sponge cake, dried fruits, and lactobacillus preparation, were replenished at approximately 16:00. The photoperiod of the colony room was 12 L:12 D (light period: 8:00-20:00, dark period: 20:00-8:00). The observations and experiments were conducted in the animals’ home cages between 8:00 and 17:00. All marmosets were well habituated to the presence of the experimenters (K.S., S.Y.-N., and a technical staff member) in the colony room to conduct the observations and experiments.

### Evaluation of the caregiver-infant relationship

Caregiver-infant interactions were studied through four structured assays of parental behaviors as described ^38^: briefly, (1) instantaneous scan sampling of the family cage, 5 times per day from PND 0 to PND 92 (minimum and maximum data points in the dataset; the same hereinafter), to record the identity of the carrier(s) and the number of infants being carried; (2) continuous 20-min focal observation of the family from PND 0 to 60, to record each family member’s caretaking behavior, social behavior, and nonsocial behavior in each 30-sec bin; (3) dyadic “infant-retrieval assay”, starting from 1 to PND 41; and (4) one-to-one “food transfer assay” from PND 69 to PND 128. In our previous study, all of these data were coded on-site for caregiving behaviors and used to yield a set of parameters as summarized in Table S3 (twins are pooled because the individual was not distinguished in the observation) ^38^. From this dataset, we performed microanalyses of infant retrieval assays for both caregivers and infants as described below.

### Infant retrieval assay

The dyadic infant retrieval assays, or brief separation-reunion sessions from the infant’s viewpoint, were based on the previous marmoset literature ^38,66^. The stress loaded onto animals was minimized using their home cage as the testing arena, although some stress was inevitable because of the temporary separation of other family members from the testing area and a brief period (maximum 600 sec) of infant isolation.

The infant retrieval assay (Fig. 1A) was conducted from PND 1 to 41 as described ^38^. An infant was presented to one of its parents or elder siblings when we returned an infant to its home cage after daily body weight measurement. If more than two caregivers from each family were tested in the same parturition or an infant was a singleton, the infants were separated twice on the same day. The order of the test of multiple family members was counterbalanced. When there were twins in one family, the stimulus infants were alternated. All family members were acclimatized to the tunnel and the wire mesh basket without infants before the test. During the test, the joined home cages were divided into three parts (43 × 66 × 60 cm each) using steel partitions. Two of these cages were connected by a mesh tunnel (75 × 30.5 × 21 cm) and used for the test (Fig. 1A). The caregiver was placed in the left cage before the start of the test. The other family members were placed in the third cage not used for the experiment. During this procedure, the caregiver and the other family members were gently separated by being lured with a piece of sponge cake to minimize the effect of handling on the caregiver’s subsequent behavior. Then, the infant was gently taken up from a carrier and placed into the mesh basket (15 cm in diameter × 15 cm high), which contained a gauze-covered electric hand warmer (KIR-SE1S, Sanyo, Osaka, Japan) to maintain the body temperature of the infant during separation. The infant in the mesh basket was placed in the right cage. Opening the shutter of the caregiver’s cage permitted the caregiver to access the infant’s cage. The behavior of the caregivers and infants before retrieval and 600 sec after retrieval, or for 600 sec after the opening of the shutter when retrieval was not attempted, was directly observed and recorded using two video cameras (HDR-AS100V, Sony, Tokyo, Japan) as well as a directional microphone (MKH 416, Sennheiser, Hanover, Germany) connected to a linear PCM recorder (DR-60DMKII, Tascam, Tokyo, Japan). The audio was recorded at 24-bit and 96 kHz. The time from the opening of the sliding door to the retrieval of the infant, which was defined as when all of the infant’s limbs were in contact with the caregiver’s body, was recorded as the retrieval latency. Immediately after successful retrieval, the caregivers’ infant-directed behaviors were coded on-site for 600 sec with 30-sec bins. The session ended if the caregiver did not retrieve the infant for 600 sec (or 300 sec for the initial 12.6% of these experiments). Twenty-three (2.8%) and 39 (4.8%) sessions ended without retrieval for 300 sec and 600 sec, respectively, among 815 sessions in total.

### Microanalysis of the infant retrieval assay

Using infant retrieval assays recorded by video and a directional microphone ^38^, manual microanalyses were completed for 286 sessions using 7 families (Table S1) and investigated in this study. The video was analyzed with 0.2-sec bins for the behaviors listed in Table S2 using Solomon Coder (ver. 19.08.02). Infant’s and caregiver’s vocalizations were detected and classified from the spectrogram of the audio using PRAAT (ver. 6.0.43). For the detailed vocal analyses, we used data from 171 sessions with high-quality vocal recordings.

In the statistical analyses, 21 sessions during the first postnatal week with the caregiver who performed the infant retrieval assay for the first time were regarded as training and excluded from the statistical analysis ^38^. The percentage of rejection (*%Rejection*) and infant calls during carrying were calculated in a session with at least one retrieval. The dyadic averages of these parameters were not calculated in an infant-caregiver dyad, Pearl-Mogol, because Mogol did never retrieve Pearl in the three sessions used in the analysis.

Among 164 observed dismounts, 34 dismounts occurred without a preceding rejection. The other 130 dismounts occurred after the rejection with variable intervals ranging from 0.4 sec to 417.2 sec. To divide these events into forced and voluntary dismounts, we performed log-survivorship analysis for rejection-dismount intervals; the 130 dismounts that occurred after a rejection are plotted for the intervals from the preceding rejection in a serial order. Then, to divide these events into forced and voluntary dismounts, we performed a segmented regression analysis and found the inflection point at 9.37 sec. The rejection-dismount intervals shorter than this threshold were regarded as influenced by the preceding rejection (66.5%, 109 out of 164 dismounts). Twenty-one dismounts with an interval less than the threshold and 34 dismounts without any preceding rejection were regarded as voluntary dismounts (33.5%). Voluntary dismounts were rare for infants younger than PND 28 (23.4%, 25 out of 107 cases), while 30 out of 57 cases and 52.6% of dismounts were voluntary on or after PND 28.

To examine the selectivity of infant attachment, we conducted an additional seven sessions using three infants and two unfamiliar, biologically unrelated adults (one male, Oji, and one female, Hime) with multiple parental experiences. The unfamiliar adults were employed for the experiment when they did not nurture preweaning infants. To minimize the chance of aggressive attacks toward the infants by unfamiliar adults, we selected the adults with comparatively less rejecting as caregivers and monitored their behavior on-site to be able to interrupt the experiments if the adults attacked the infants. In the present study, there was no aggression by the adults exceeding the level of normal rejections. To evaluate the caregivers’ behavior, seven sessions with two of their own infants were compared as controls (own infant). For the control of the infants’ behavior (own parent), seven sessions with two mothers and two fathers of the infants were used. The sex of caregivers and age of infants in the control sessions were matched.

### Call annotations

Infant and caregiver calls and call types were annotated manually. As previously reported ^49,67^, infant and adult calls are distinct spectral and temporal characteristics; infant phees are shorter and higher in frequency (Fig. S5B-C); each twitter phrase is associated with a downward frequency modulation at the end (“twitter-hook”) (Fig. S5B-D). The experimenter identified the characteristic calls produced by the infant or the caregiver when the calls were clearly associated with body movements or mouth opening of either individual using two cameras (Fig. S5 and Supplementary movie 8), then inferred the call identity with this spectrographic pattern for the calls when the body or mouth of the caller was not clearly visible. With a directional microphone, the calls from other cages (inevitable as the rest of the family members were contained in the next cage during the retrieval assay sessions) appeared blurred and low intensity in the spectrogram (Fig. 2E bottom, the phee and twitter calls around the timing of retrieval. Compare the following twitter call by the caregiver). When the source of the call could not have been confidently identified, such an ambiguous call was omitted from the analysis. Using this strategy, the infant calls exhibited suitable interrater reliability (Cohen’s к = 0.844 (Vhee), 0.527 (Phee), 0.742 (Trill), 0.835 (Tsik) and 0.813 (Twitter), where the chance level к = 0). For the caregivers’ vocalizations, we spared detailed analyses for the later study, because our screening analysis in Fig. 1c suggests that the caregivers’ physical behaviors such as retrieval and rejection are more immediately influencial than caregivers’ vocalizations on infant behaviors and calls during the free dyadic interactions (see Fig. S5A), and the main scope of this study is to identify the caregiving influences shaping infant attachment behaviors.

### Artificially reared infants

In our breeding colony, we could often maintain the whole litter of triplets with supplemental feeding but not always, depending on the capacity of milk production of the mother and the total caregiving motivation of the family members. We utilized 5 infants, each out of such triplet cases (or a twin in the case of Chuck’s family, as shown above). The details of rearing conditions are described for individual cases in Table S6. Briefly, infants were isolated from their families on the day of birth. During isolation, they were kept individually in small cages (22 × 14 × 14 cm or larger according to their growth), given a rolled soft cloth to cling to, and placed in an electric incubator to keep the ambient temperature at 30–37°C (adjusted to their development). The incubator was placed in the same colony room except for two infants (Senazo and Senako), but not necessarily close to the family cage, and visual, olfactory, and auditory communications with other marmosets were constrained but not completely eliminated. During the daytime, each infant was reunited with the family 2-6 hr/day, 3-7 times per week, except for one infant (Michal) that had received aggressive rejections from the family members. Experimenters and care staff handled these Art infants for milk feeding (3 times /day), body weight measurements (once/day), and for experiments. Human baby food, chow soaked in milk, and normal chow after 1.5 months old were also provided. After weaning, each Art infant was put into a normal home cage (42 or 43 (wide) × 66 (high) × 60 (deep) cm) with normal chow and water (ad libitum) and singly housed in the same colony room with their family. The body weight of artificially reared infants tended to be lighter than that of control littermates (*p* = 0.0792), but it was within the range of variations of the family-reared infants (Fig. S6).

The family-reared littermates of the artificially reared subjects were used as control subjects, except for Atako. Because Atako’s original family was assigned to another experiment, Atako’s experiments were conducted with the Chuck family, which bore two infants one day before Atako’s birth. Therefore, Atako’s age-matched control was assigned to one infant of Chuck’s family. Atako died in an accident at PND27, so no further experiments were conducted.

### Family reunion observations of artificially reared infants

Four Art and five Cont infants (Table S7) were repeatedly employed for the family reunion observations during PND 1–53. The observations were performed once to three times per week. The observations were conducted in a home cage (size is described in Table S7) that contained a mother, a father, and one to three older siblings. First, from the family cage, the family-reared Cont infants were removed and subjected to body-weight measurements. Then, Art and Cont infants were introduced into the family cage in random order. We used the period of their stay in the home cage from the introduction and without interruption by daily cage cleaning for quantitative behavioral analysis, of which length varied between 16 and 119 min (mean ± standard deviation: 57.97 ± 25.93 min), as it turned out that the cleaning significantly altered their behavior.

The caregivers’ reaction when the infant was returned to their home cage was categorized as follows: approached, the caregiver approached the cage entrance where the infant was released by the observer; and helped, the caregiver helped the infant to hold its body or tried to attack the observer’s hand holding the infant.

The following infants’ conditions and behaviors were manually categorized using *one-zero sampling* in 10 sec bins: condition - carried, rejected, attacked, and touched by family members; behaviors - calling. The infants’ calling was determined by the sound and oral and/or abdominal movements of the infant using video. When the source of the vocalizations could not be attributed to one infant, because multiple individuals were calling or their movements were unclear, it was recorded as “calls by unspecified infant(s)” and excluded from the analysis. The avoidant behaviors of the infants were observed continuously.

### Infant retrieval assay with artificially reared infants

Artificially reared infants and their control subjects (artificially reared: n = 4, control: n = 5, PND 2–30, 69 sessions) were repeatedly employed for the retrieval assays in the home cages of their families (Table S8). It was performed twice per week. The detailed procedure is described above. The latency of entering the infant cage, touching the infant, and retrieving the infant was measured. The behaviors of caregivers and infants were analyzed in 0.2 sec bins as shown above. The caregiver’s body part where the infant was holding was determined when the infant stopped just after a carrying bout started.

### Vocal recordings in isolation

Ten infants (artificially reared male: n = 1 (Michael), control: n = 9 (four males (Cubby, an infant from Chuck’s family, and two infants from Oji’s family. The three males were not used in the other experiments of the present report), and five females (Eugenie, Shirayuri, Suisen, Mimosa, Kodemari), 170 sessions, PND 3–104) were used for the vocal recordings in isolation. The recordings were performed once or twice per week. It was conducted in another room separated from the colony rooms. The subject was removed from the home cage and transferred to a plastic cage (28 cm × 44 cm × 20.5 cm) placed in the recording room. The experimenter left the room and recorded the infants’ vocalizations using a video recorder and the sound recorder (Sennheiser MKH416-P48U3, TASCAM DR-60DMK II) for 600 sec. The vocalizations were manually analyzed on PRAAT.

### Statistical analysis

Statistical analyses were conducted using R software (version 3.6.3) ^68^. In the retrieval assays with the normative family, statistical analyses were performed by the generalized linear mixed model (GLMM) with some exceptions. The glmer function of the lme4 package ^69^ was used for the GLMM. The infant and the carrier were added as random effects with an intercept to control pseudoreplication. The fixed effects included in the models are mentioned in the figure legends. The best models were selected using the dredge function in the MuMIn package ^69^. For the correlation matrix, we conducted Pearson’s product-moment correlation analysis. Correlation coefficients and *p value*s adjusted with Holm’s method or Benjamini_JHochberg’s method were calculated using the corr.test function in the psych package ^70^. For the segmented relationships in regression models, we employed the segmented function in the segmented package ^71^. The difference in filial behavior was analyzed using the Wilcoxon rank sum test. The Wilcoxon signed-rank test with continuity correction was conducted for the comparison of infant calls for 10 sec after and before the shift of the social contexts.

In the family-reunion observations and the retrieval assays using artificially reared marmosets, statistical analyses were performed by GLMM. The family and the infant nested in the family were added as random effects for family-reunion observations, and the caregiver and the infant were added as random effects for retrieval assays to control pseudoreplication. Random effects included intercepts. PND, artificial rearing, and their interaction were added as a fixed effect in the initial model. The best models were extracted using the dredge function in the MuMIn package ^72^. For count data with/without an upper limit, binomial/Poisson distributions were adopted. Latency data were transformed logarithmically and then treated as Gaussian distributions. Other data were treated with normal distributions. For the analyses of the holding site in the retrieval assays, the clmm function in the ordinal package was employed instead of the glmer function in the lme4 package to deal with the ordinal response variable ^73^.

### Data and materials availability

All data needed to evaluate the conclusions in the paper are present in the paper and/or the Supplementary Materials. The raw video and audio files can be provided by K.O.K. pending scientific review and a completed material transfer agreement. Requests for the these raw data should be submitted to.

## Supplementary Information

**Figure S1 Inter-observational reliability between the on-site coarse behavioral coding and off-site, detailed behavioral microanalyses of the infant retrieval assay.**

The on-site analyses performed with 30-sec bins (by rater S.K. (38)) are compared with the off-site analyses with 0.2-sec bins in this study (by rater A.T.), by converting A.T. ‘s coding results into the 30-sec bins and classifying them as “carried” (blue), “rejected” (yellow), or “not carried” (red). The interrater reliability between the two raters was assessed by Cohen’s kappa (κ = 0.926, p < 0.001).

**Figure S2 Infant behaviors in the retrieval assay using family-reared infants.**

A Segmented regression analysis of the rejection-call interval. The inflection point could not be determined; thus, we also used the 9.4-second offset obtained from movement analysis (Fig. 2B) for call analysis. A total of 77.18% (3981 out of 5158) of the infant calls during carrying occurred during and within 9.4 sec after the preceding rejection.

B Spectrogram of the infant’s vhee call.

C Developmental changes in infant call frequencies in each social context. The columns and bars indicate the mean ± s.e. of total calls. The color shows the type of call. This graph corresponds with Fig. 2G. Only the data with detailed vocal analysis were included. The numbers within the parentheses are the numbers of the sessions.

D Scatter plots and regression lines of infant calls during carrying (top) and avoidant behavior (bottom) and caregiving parameters (left: Ret_latency, center: %Rejection, right: %Carrying). More avoidant behaviors were observed in the dyad with longer retrieval latency (t = 2.76, p = 0.0078), although the infant calls during carrying were not different from those in the other dyad. In the dyad with a higher %Rejection, the infant calls during carrying and the avoidant behaviors were more frequent (call-during-carrying: t = 5.13, p < 0.001, avoidant behavior: t = 4.323, p < 0.001). In the dyad with a higher %Carrying than the average, the infant calls during carrying and the avoidant behaviors were less frequent (call-during-carrying: t = −4.25, p < 0.001, avoidant behavior: t = −7.367, p < 0.001). Each dot shows the value of each session, and the same color indicates data from the same caregiver.

E Violin plots of the infant calls during holding (top) and transport (bottom) in the groups of dyads divided by high/low parental behaviors (retrieval latency, %Rejection, %Carrying). Each dot shows the value of each session, and the color indicates the individual caregiver. The filled circles and error bars show the mean ± s.e. The numbers within parentheses are the numbers of the sessions.

GLMMs were used for statistical analyses. **: p < 0.01, ***: p < 0.001

**Figure S3 Vocal behaviors in isolation.**

A-F The mean ± s.e. call frequencies in the isolated recordings (Art: n = 1 male; Cont: n = 9 (4 males, 5 females). Total calls (A) as well as phee (B), vhee (C), tsik (D), trill (E), and twitter calls (F).

G The mean ± s.e. total calls and composition of call types of Cont (left) and Art infants (right). Error bars represent the standard error of the total calls. The left is the same as in Fig. 2I.

**Figure S4 Family reunion observation with artificially reared infants.**

The duration of a carrying bout (A), the latency of the first rejection (B), and the number of caregivers’ rejection bins (C) in a carrying bout were calculated. A carrying bout is a period from the bin when an infant starts to cling onto a caregiver to the bin when the infant dismounts from the caregiver or moves to another caregiver. The shape and color of the markers represent each infant. Top: family-reared infants (Cont), bottom: artificially reared infants (Art).

A The duration of a carrying bout (Cont: 363 bouts, Art: 299 bouts) was shorter among the older infants (t = −5.63, p < 0.001).

B The latency of the first rejection in a carrying bout (Cont: 323 bouts, Art: 244 bouts) was shorter among the older infants (t = −5.10, p < 0.001).

C In the number of caregiver rejection bins (Cont: 363 bouts, Art: 299 bouts), the interaction effect of Art and PND was significant (z = −7.65, p < 0.001), indicating that the coefficient of PND among the artificially reared infants was smaller than that among the family-reared infants.

Statistical analyses were performed by the generalized linear mixed model (GLMM). *: p < 0.05, **: p < 0.01, ***: p < 0.001

**Figure S5 Infant and caregiver calls in the infant retrieval assay.**

A Raster plot of a representative retrieval assay session participated by John at PND19 and its father Chuck. Infant’s Moving alone, Searching for contact, caregiver’s Breast feeding, Licking, Grooming, Running, Stereotypy, Self-grooming, Marking, Feeding, and Yawning are not shown in the plot because these behaviors were never performed in this session. The dashed square indicates the part shown in Supplementary movie 8. Note that the infant called vigorously before the first retrieval and did not call afterward, until the paternal rejection started. At the session start, the father exhibited an immediate physical approach toward the vocalizing infant but did not vocally respond to the infant calls. The father emitted twitter and phee during carrying without rejection, presumably directing to other family members in the adjacent cage (see C), while the carried infant remained silent.

B The spectrograms of infant calls. Top: the raster plot of infant call types presented by the middle spectrogram. *: calls confirmed by the infant’s body movements (see Supplementary movie 8). Bottom: Magnified spectrograms of infant-typical call characteristics. Compare the length of phee with the father’s phee (C).

C Example spectrograms of caregiver calls. *: calls confirmed by the mouth opening (see Supplementary movie 8). These father’s calls were interspersed by twitter calls from the other cage.

D An example spectrogram of infant twitter (George PND 9, Fig. 2E bottom).

**Figure S6 Body weight of artificially reared and family-reared infants.**

A Infant body weight and LOESS fitted curve. Warm colors: artificially reared infants, cold colors: control littermates.

B Estimated body weight at PND40 of the artificially reared infants and control littermates. The body weight of artificially reared infants tended to be lighter than that of control littermates, although it was not at a significant level.

C Estimated body weight at PND40 of the artificially reared infants and all family-reared infants. The body weight of artificially reared infants was within the range of variations of the family-reared infants.

**Table S1 List of family-reared infants and their retrieval assay sessions.**

The numerals indicate the postnatal day of the infant in the session. Parentheses indicate the first week sessions performed by the caregiver without previous experience of infant retrieval assays and were excluded from the statistical analysis. Italics: indicate the sessions without high-quality vocal recordings, which were excluded from the detailed vocal analyses. Underlines indicate the initial 16 sessions that were censored at 300 sec when the first retrieval did not take place, while the others were censored at 600 sec. Red: female, blue: male. See also (38).

**Table S2 List of observed behaviors in the retrieval assay.**

**Table S3 List of parameters employed in our previous paper.**

These parameters were obtained in (38) and were used for the correlation matrix in Fig. 1C.

**Table S4 The r values of the correlation matrix in Fig. 1C**.

Red and blue indicate positive and negative correlations, respectively.

**Table S5 The p values of the correlation matrix in Fig. 1C**.

Red: p < 0.05.

**Table S6 List of artificially reared infants.**

**Table S7 List of family reunion observations with Art infants.**

Cont: Control, Art: Artificially reared.

**Table S8 List of retrieval assays with Art infants.**

Cont: Control, Art: Artificially reared. “Cage size” represents the size of a home cage. In the test, two home cages were connected by the tunnel.

**Supplementary movie 1 Typical retrieval in the retrieval assay**

Wendy (PND 15) was left alone and called severely, and then Fastener (mother) retrieved Wendy. After the retrieval, Wendy stopped calling.

**Supplementary movie 2 Caregiver’s rejection in the retrieval assay**

Gaku (PND 13) was carried and then rejected by Tochan (father). After the rejection, Gaku dismounted and called intensely while alone.

**Supplementary movie 3 Direct transfer of family reunion observation**

Yoshie (PND 1) was rejected by Junior (elder brother) and called severely. Yukita (father) approached them (0:18) and then received Yoshie from Junior (0:23). After Yukita’s retrieval, Yoshie stopped calling.

**Supplementary movie 4 Dismounting of infants during carrying in family reunion observation**

The Art infant Senazo (PND 23) clung to the father (Yukita) (0:14) and soon dismounted without distinct rejection (0:23). Then, the Cont littermate Yoshie clung to Yukita (0:32) and was rejected (0:40∼). Yoshie kept clinging during the rejection for more than 50 sec and finally dismounted (not included in this video).

**Supplementary movie 5 Avoidant behavior of an artificially reared infant in the retrieval assay.**

The Art infant Michael (PND 15) clung to the trunk of the father Chuck and dismounted without distinct rejection (0:19). Chuck tried to retrieve Michael again (0:31), but Michael refused to cling. This movie is from the session at the bottom of Fig. 6B (approximately 150-190 sec).

**Supplementary movie 6 First retrieval of a control infant in the retrieval assay**

After the first retrieval, the Cont infant (Cubby, the littermate of Michael, PND 15) quickly moved to the back of the father (Chuck).

**Supplementary movie 7 First retrieval of an artificially reared infant in the retrieval assay**

After the first retrieval, the Art infant Michael (PND 15) moved onto the body of the father (Chuck) (0:42∼). Michael occasionally clung to Chuck’s head and face, which induced rejection by Chuck (0:55∼). Michael took a longer time to settle to a suitable position than Cubby.

**Supplementary movie 8 Call annotations in the retrieval assay**

The infant John (PND 19) was retrieved by the father Chuck. See also Figure S5.

