## Supplementary figures and images for "Anxious about rejection, avoidant of neglect: Infant marmosets tune their attachment based on individual caregiver’s parenting style"

### Supplemental Figure 1-6

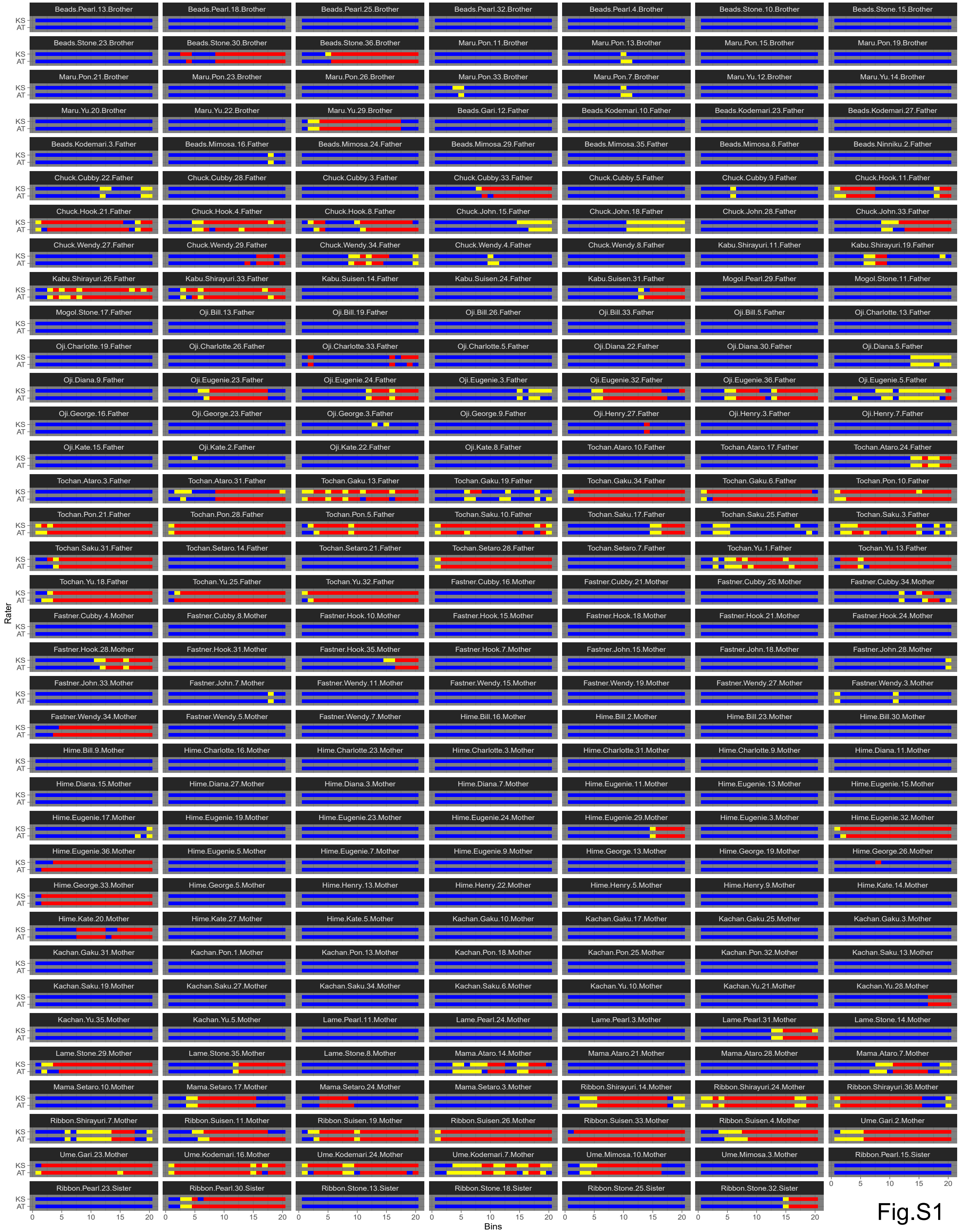

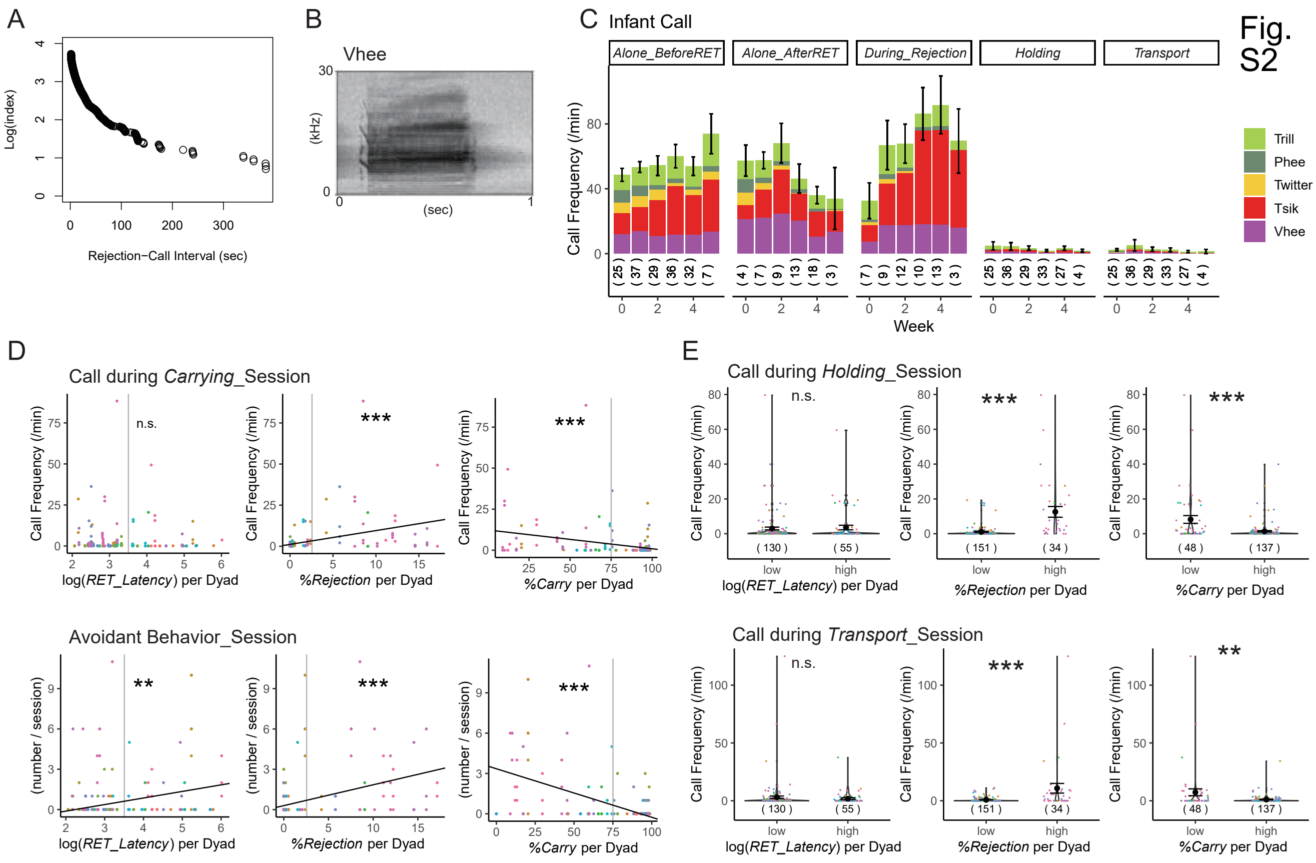

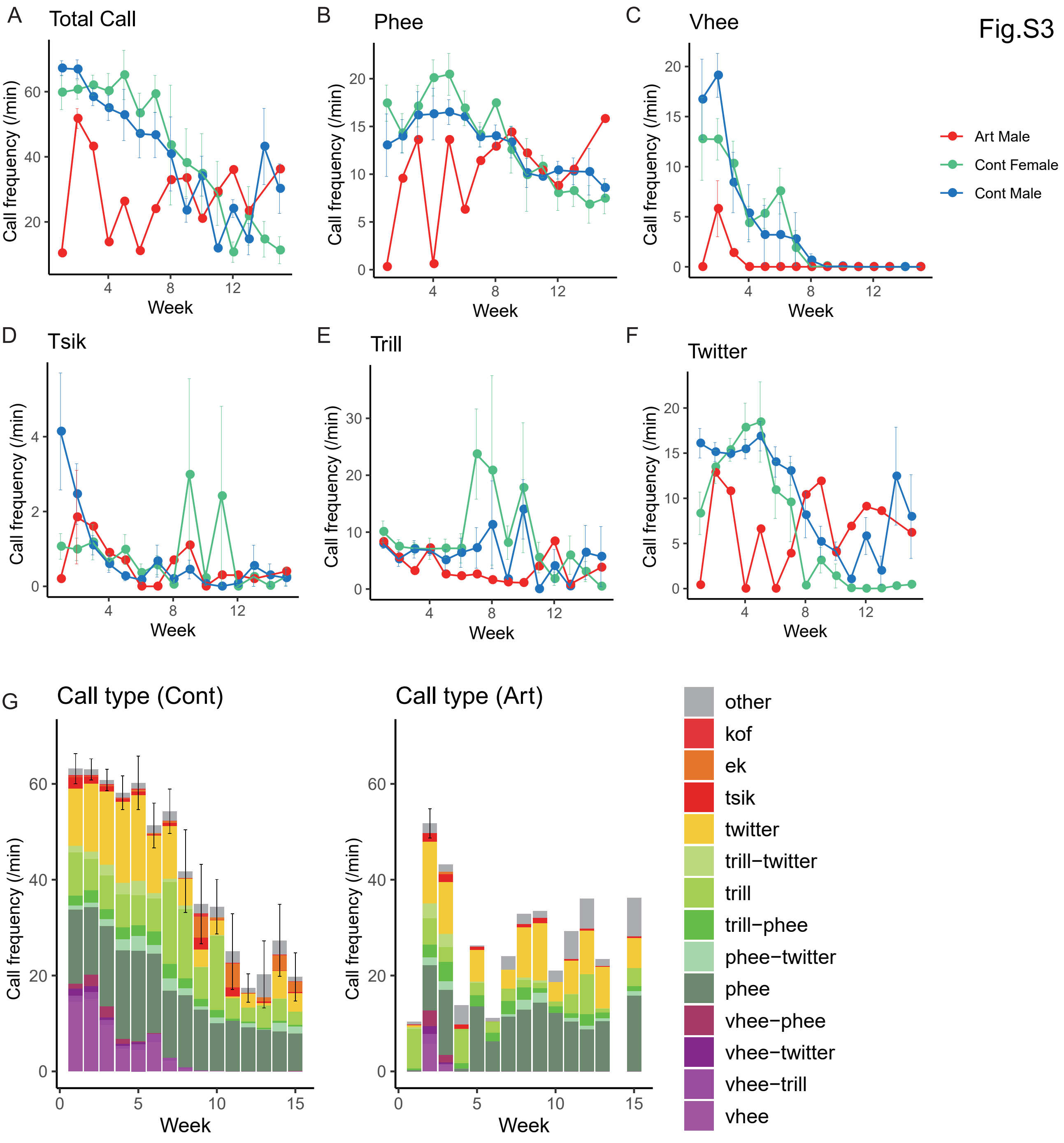

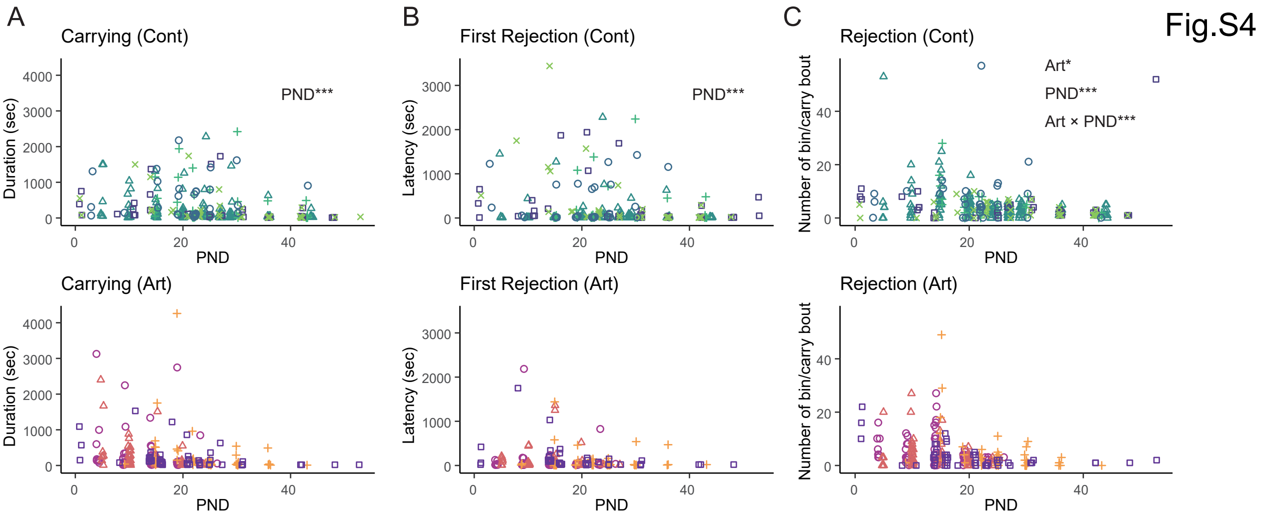

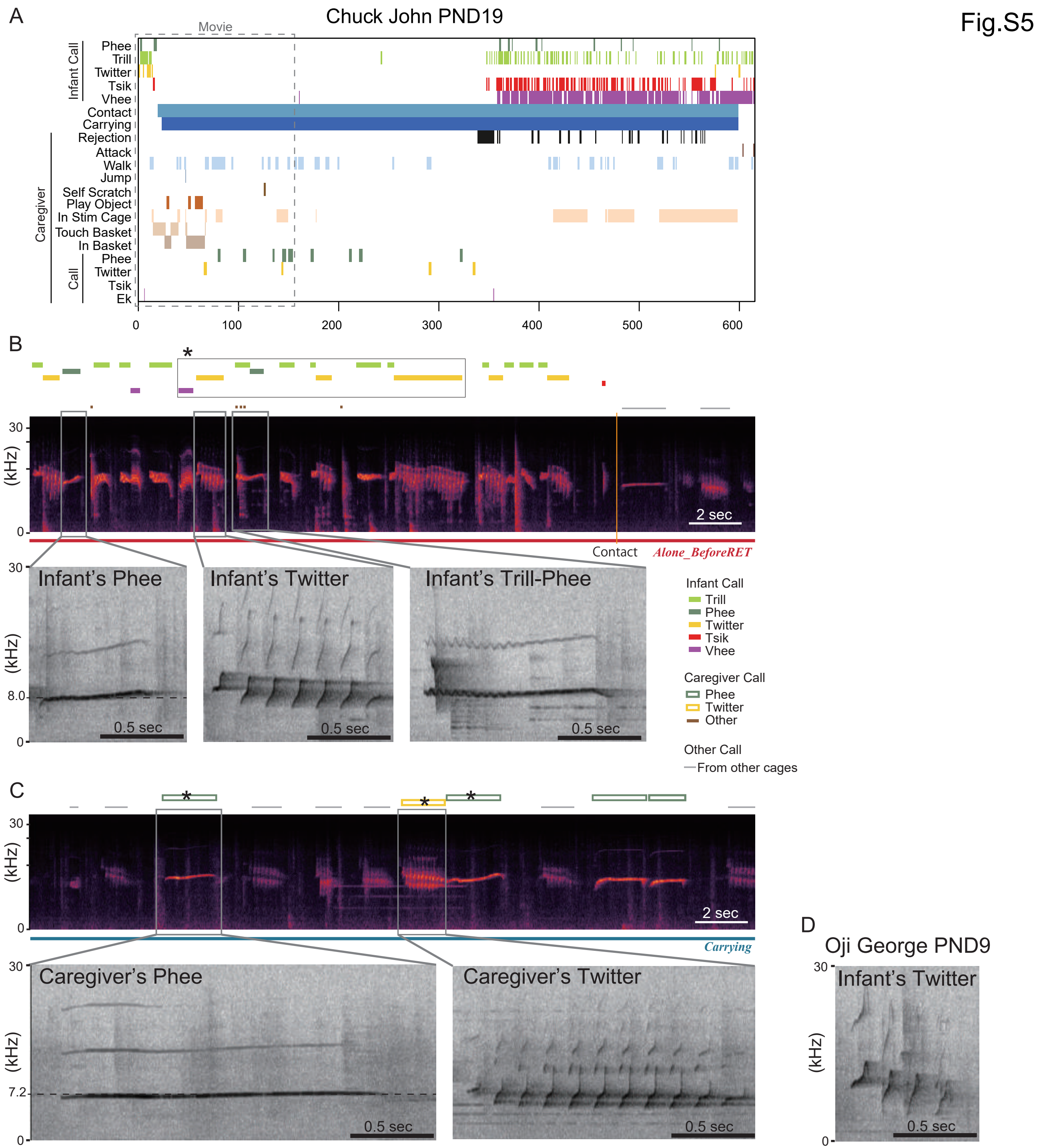

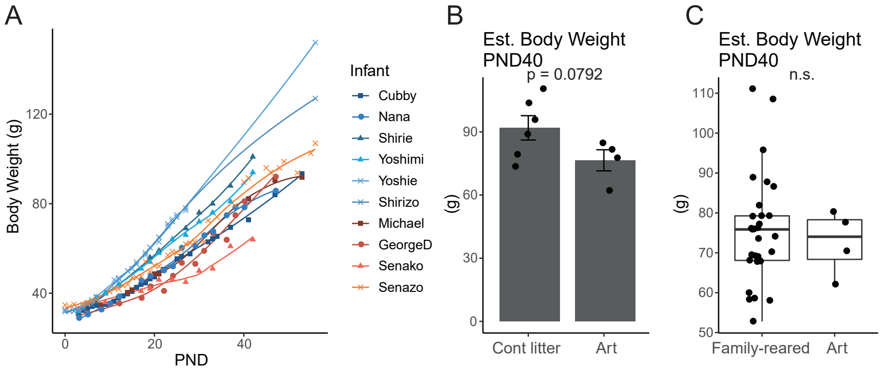
